# Characterization of [^3^H]AZ12464237 as a high affinity, non-nucleotide antagonist radioligand for the P2Y12 receptor

**DOI:** 10.1101/2024.10.10.617599

**Authors:** E. Johanna L. Stéen, Efthalia-Natalia Tousiaki, Lee Kingston, Berend van der Wildt, Nikolett Lénárt, Wissam Beaino, Mariska Verlaan, Barbara Zarzycka, Bastian Zinnhardt, Ádám Dénes, Luca Gobbi, Iwan J.P. de Esch, Charles S. Elmore, Albert D. Windhorst, Michael Honer, Rob Leurs

## Abstract

The purinergic receptor P2Y_12_ (P2Y_12_R) is a well-recognized target for anti-thrombotic agents. This receptor is also expressed in microglia, where it plays a key role in neuroinflammation and microglia activation. To investigate P2Y_12_R-mediated actions in the central nervous system (CNS), the development of novel brain-penetrant ligands is essential, along with further *in vitro* studies. A radiolabeled, easily accessible tool compound would significantly advance such drug discovery efforts. Herein, we describe the ^3^H-labeling of a non-nucleotide P2Y_12_R antagonist AZ12464237, and its *in vitro* binding properties to the receptor in membrane preparations form transfected cells, as well as on mouse brain tissues. The radioligand shows high affinity toward both the human and rat P2Y_12_R in transfected cells, with *K_d_* values of 3.12 ± 0.70 nM (human) and 16.6 ± 3.4 nM (rat), as determined by saturation binding studies. The binding kinetics of [^3^H]AZ12464237 are rapid with a short target residence time (∼1 min). We further confirmed the selectivity of the radioligand by performing competitive displacement studies, in which reported P2Y_12_R ligands and other P2Y receptors ligands were tested for binding against [^3^H]AZ12464237. Additionally, the radioligand proved useful for *in vitro* autoradiography studies on mouse brain tissues, although a small amount of off-target binding was observed in P2Y_12_R knock-out mice. This could be traced to glycogen synthase kinase 3 α. Considering the growing interest in P2Y_12_R as a biomarker for microglia activation, [^3^H]AZ12464237 represents a promising tool for *in vitro* studies, including screening assays aimed at identifying novel P2Y_12_R ligands for CNS applications.

## 1. Introduction

The P2Y receptors compose a subfamily of eight G protein-coupled receptors (GPCRs) belonging to the class of rhodopsin-like receptors, also known as family A GPCRs. The P2Y receptor subfamily can be further divided into two different subgroups based, among other things, on which G protein(s) they couple to. The P2Y_1_, P2Y_2_, P2Y_4_, P2Y_6_ and P2Y_11_ receptors in the first subgroup couple predominantly to members of the G*_q/11_* proteins, whereas the second group, including P2Y_12_, P2Y_13_ and P2Y_14_ receptors signals primarily through the G*_i_* family. In addition, the endogenous agonists differ amongst the receptors.(1,2) The P2Y_1_R, P2Y_12_R and P2Y_13_R are activated by adenosine diphosphate (ADP), the P2Y_11_R by adenosine triphosphate (ATP), the P2Y_2_R by ATP and uridine triphosphate (UTP), the P2Y_4_R by UTP, the P2Y_6_R by uridine diphosphate (UDP), and the P2Y_14_R is activated by UDP and UDP-sugar derivatives (UDP-galactose and UDP-glucose).(3)

The P2Y receptors have a wide tissue distribution and are involved in numerous of physiological responses and pathophysiological processes. As a consequence, they are highlighted as biomarkers and potential drug targets for a variety of conditions such as, dry eye syndrome, inflammation and pain, and cardiovascular diseases.(1) A key example in this family is the P2Y_12_R, which is highly expressed on platelets and mediates *e.g.*, ADP-induced platelet aggregation. Over the years, numerous structurally diverse ligands acting as P2Y_12_R antagonists have been developed, and quite a few have become important drugs in antiplatelet therapy. Cangrelor (Kengreal^®^), ticagrelor (Brilinta^®^), prasugrel (Effient^®^) and clopidogrel (Plavix^®^) are just a few examples of clinically approved drugs. The two latter, prasugrel and clopidogrel, are prodrugs based on a thienopyridine scaffold. After *in vivo* activation, their active metabolites are suggested to irreversibly bind to cysteine 97 (Cys97), located in an allosteric binding site.(4) More detailed structural information regarding binding modes was obtained in 2014 when three crystal structures of the receptor were solved. The obtained structures had the receptor in complex with full agonist 2-methylthio- adenosine-5’-diphosphate (2MeSADP), partial agonist 2-methylthio-adenosine-5’-triphosphate (5MeSATP) and non-nucleotide antagonist AZD1283, respectively. There is a striking difference in receptor conformation when bound to the agonists (but in the absence of G-protein) compared to the AZD1283-bound. In the latter, the orthosteric binding site is exposed in an open receptor conformation, whereas in the agonist-bound structures the site is compressed due to conformational changes in the extracellular region and an inward movement of helices 6 and 7.(5,6) The last decade, molecular docking and molecular dynamics have further contributed to predicting binding modes and ligand recognition of other chemical scaffolds.(7) Additional structural insights were obtained from newly solved structures; one with inverse agonist selatogrel (X-ray), as well as a cryo-EM structure of a fully active conformation of the P2Y_12_R-2MeSADP complex in the presence of G*_i_*.(8,9)

In addition to being expressed on platelets, the P2Y_12_R is also found in the central nervous system (CNS) where it is expressed on microglia, the brain’s first line of defense upon injury or inflammation.(10–13) Microglia are highly responsive to stress and injury, initiating a highly dynamic activation process that spans over a spectrum of functional microglia states depending on the surrounding environment. A pro-inflammatory neurotoxic (M1) microglia and anti-inflammatory neuroprotective (M2) microglia represent the two extremes of the spectrum.(14) The P2Y_12_R is upregulated on M2 microglia and downregulated on M1 microglia.(13,15) An expression profile that makes this receptor a promising biomarker for the anti-inflammatory microglia state. Whereas the M1 microglia have been extensively studied by using tool compounds targeting biomarkers, such as cannabinoid receptor type 2, cyclooxygenase-2 and purinergic 2X type 7 receptor (P2X_7_R), the M2 has not been thoroughly characterized.(16) Therefore, there is a growing interest in developing a tool compound targeting the P2Y_12_R in the CNS, which would allow for studying the dynamical changes in microglia activation during onset and progression of different CNS diseases. Moreover, understanding the detailed mechanisms of P2Y_12_R-mediated functions in the brain can open the way for new treatment opportunities and support drug development. Unfortunately, the ligands developed for targeting peripheral P2Y_12_R, including the clinically approved drugs, do not have ideal physicochemical properties for targeting the receptor in the CNS. In general, they all have high topological surface areas (TPSA), high molecular weights and are charged at physiological pH.(17) Therefore, new and structurally different ligands will have to be developed and robust *in vitro* binding assays are needed to guide and facilitate this development.

So far most of the reported *in vitro* assays to study P2Y_12_R have typically relied on measuring its functional activity with respect to platelet aggregation. The reported competitive P2Y_12_R binding assays have typically made use of nucleoside-based ligands radiolabeled with either ^3^H or ^125^I, some of which are shown in Figure 1.(18–22) In the early 2000s, several high-throughput screening (HTS) campaigns by different pharmaceutical companies, resulted in the discovery of a number of novel non-nucleoside P2Y_12_R antagonists. For instance, Bach *et al*. reported a series of ethyl nicotinate derivatives as high-affinity antagonists for the P2Y_12_R. Out of these, derivative AZ12464237 showed an IC_50_ value of 6.3 nM in competitive P2Y_12_R binding studies using [^125^I]AZ11931285 (Figure 1).(23) We have previously investigated the potential of a ^11^C-labeled analog of this compound in positron emission tomography (PET) imaging of the P2Y_12_R during neuroinflammation.(24,25) Unfortunately, this compound does not enter the brain. Nonetheless, it can serve as a useful tool to study P2Y_12_R *in vitro*. Herein, we report the radiosynthesis and *in vitro* binding characteristics of the ^3^H-labeled analog of AZ12464237 ([^3^H]AZ12464237) to the P2Y_12_R on membrane preparations, cell pellets and mouse brain tissues. To further support the relevance of this compound as a tool for *in vitro* studies, we used it in competitive displacement studies with other reported P2Y ligands.

**Figure 1.**
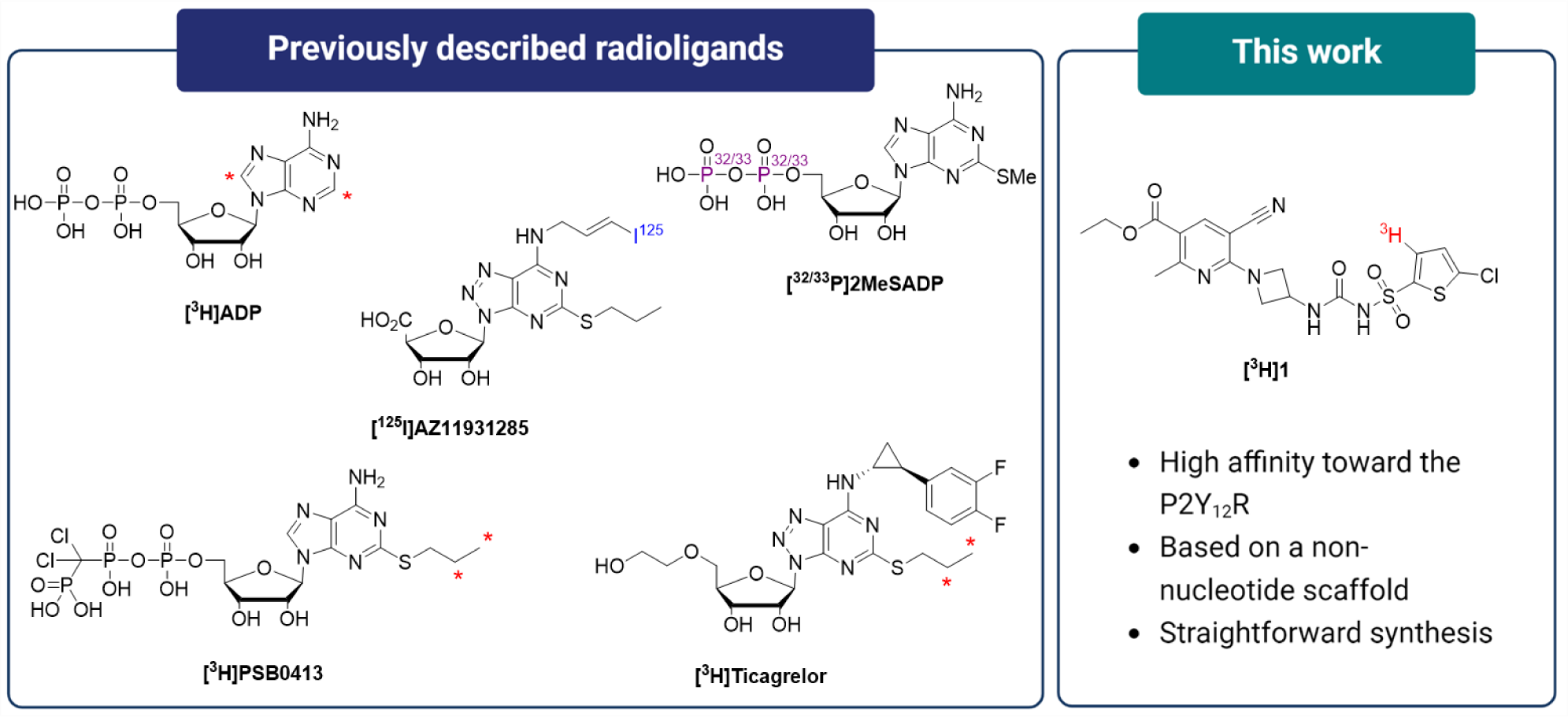
Previously described radioligands used for studying P2Y_12_R binding *in vitro* and/or in screening assays, and the radiolabeled ethyl nicotinate derivative ([^3^H]AZ12464237) characterized in this work. Red asterisks indicate ^3^H-label positions.

## 2. Materials and methods

### 2.1. Materials

Cell culture media was obtained from Westburg BV (Leusden, The Netherlands), additives to the culture media (Fetal bovine serum (FBS), *L*-glutamine, HEPES and Penicillin-Streptomycin) were obtained from Thermo Fisher Scientific (Bleiswijk, The Netherlands). Additives for binding assay buffer; bovine serum albumin (BSA), polyethylenimine (PEI), Triton X-100, Tween-20, Tween-80 were obtained from Merck Life Science NV (Amsterdam, The Netherlands). Compounds evaluated in competitive binding studies; 2MeSADP, AZD1283, elinogrel, ticagrelor, MRS221, MRS2578 and PPTN·HCl were purchased from Tocris (Bristol, United Kingdom) and used as obtained without further purification. Materials and reagents for synthesis of precursor and reference compounds were purchased from Merck Life Science NV (Amsterdam, The Netherlands).

### 2.2. Chemistry

#### 2.2.1. General information

Room temperature corresponds to a temperature interval from 18–21 °C. Reactions requiring anhydrous conditions were carried out under argon and using oven-dried glassware (152 °C). Nuclear magnetic resonance (NMR) spectra were acquired on a Bruker (Billerica, MA, USA) AVANCE 500 (500.27 MHz for ^1^H, 125.81 MHz for ^13^C and 533 MHz for ^3^H). Chemical shifts (δ) are reported in parts per million (ppm) relative to the solvent residual peak (DMSO-*d_6_*: *δ* 1H = 2.50, *δ* 13C = 39.5). The resonance multiplicity is abbreviated as: s (singlet), bs (broad singlet), d (doublet), t (triplet), q (quartet) and m (multiplet). Spectra were analyzed with the MestReNova (version 14.2.1-27684, Mestrelab Research S.L.) software. Electrospray ionization high-resolution mass spectrometry (ESI-HRMS) was carried out using a Bruker microTOF-Q with positive ion mode (capillary potential of 4500 V). Thin-layer chromatography (TLC) was carried out using either normal phase plates (silica gel 60 coated with fluorescent indicator F254s) from Merck (Darmstadt, Germany). Spots were visualized by ultraviolet light at 254 nm and by KMnO_4_ staining. Flash column chromatography was performed using a Büchi Sepacore Flash System (2 x Büchi Pump Module C-605, Büchi Pump Manager C-615, Büchi UV Photometer C-635, Büchi Fraction Collector C-660) with pre-packed columns. Analytical high-performance liquid chromatography (HPLC) was performed on two different systems: (1) on a Waters Acquity HPLC system with photodioarray UV detection with in-line detection of radioactivity using a lab Logic BetaRam model 5. Chromatograms were analyzed using the Laura software (version 4.2.10.111, LabLogic); (2) on an Agilent 1200 series system (Agilent, Santa Clara, CA, US) connected to a Jasco PU-2080 Plus Intelligent HPLC pump (Jasco, de Meern, The Netherlands), a Jasco UV-2075 Plus UV/vis detector, and a LabLogic radio-HPLC detector β-ram Model 4 (LabLogic Systems Ltd, Sheffield, UK). Chromatograms were analyzed using the Laura software (version 4.1.12.89); LC/MS was performed on a Agilent 1100 series on a Xselect CSH C18 3.5 μ (4.6 × 100 mm) column (Waters, Milford, MA, USA) using a gradient of MeCN (solvent B) in 0.2% formic acid in water pH 3 (solvent A) as eluent. Gradient settings: 0 → 10 min (5% → 95% solvent B); 10 → 12.5 min (95% solvent B) with UV detection at 254 nm and mass detection using a 3100 mass detector (140–780 Da). The molar activity (A*_m_*) was determined by LC/MS using Isopat2 to deconvolute the MS signals.(26)

#### 2.2.2. Ethyl 6-(3-(3-((5-chlorothiophen-2-yl)sulfonyl)ureido)azetidin-1-yl)-5-cyano-2- methylnicotinate (AZ12464237)

Reference AZ12464237 was synthesized as described by Bach *et al*.(23)

#### 2.2.3. 5-Chloro-3-iodothiophene-2-sulfonamide (1)

*N*-Iodosuccinimide (342 mg, 1.52 mmol) was added to an ice-cold solution of 5-chlorothiophene-2- sulfonamide (300 mg, 1.52 mmol) in triflic acid (700 µL, 7.91 mmol). The mixture was stirred at room temperature overnight (22 h) before it was treated with water and aqueous Na_2_CO_3_ (0.5 M), and extracted with dichloromethane (DCM). The combined organic phases were washed with brine, dried over Na_2_SO_4_, filtered and concentrated under reduced pressure. The crude product was purified by flash column chromatography on silica gel using 40% EtOAC in hexane as eluent to afford **1** (282 mg, 57%) as an off-white solid. ^1^H NMR (500 MHz, DMSO-*d_6_*) *δ* 7.89 (s, 2H), 7.50 (s, 1H). ^13^C NMR (126 MHz, DMSO-*d_6_*) *δ* 145.5, 136.0, 134.2, 85.3. ESI-HRMS: calculated for C_4_H_3_ClINNaO_2_S_2_ [M+Na]^+^, 345.8231; found, 345.8235.

#### 2.2.4. Ethyl 6-(3-(3-((5-chloro-3-iodothiophen-2-yl)sulfonyl)ureido)azetidin-1-yl)-5-cyano-2-methylnicotinate (3)

Building block **2** was synthesized according to a literature procedure.(23) A solution of **2** (27.0 mg, 0.10 mmol) and *N*,*N*-diisopropylethylamine (DIPEA; 37.0 µL, 0.21 mmol) in anhydrous DCM (1 mL) was added to triphosgene (11.0 mg, 0.04 mmol) in DCM (300 µL). The mixture was stirred at room temperature for 50 min. Sulfonamide **1** (49.0 mg, 0.15 mmol) and DIPEA (27.0 µL, 0.16 mmol) in dry DCM (200 µL) was added. The mixture was stirred at room temperature for 5 h under argon before it was diluted with DCM, washed with HCl (0.5 M) and brine, dried over Na_2_SO_4_, filtered and concentrated under reduced pressure. The crude product was purified by flash column chromatography on silica gel using a gradient of 0% → 100% EtOAc in hexane to afford **3** (40 mg, 63%) as a white solid. ^1^H NMR (500 MHz, DMSO-*d_6_*) *δ* 8.29 (s, 1H), 7.68 (s, 1H), 7.48 (bs, 1H), 4.65–4.44 (m, 3H), 4.23 (q, *J* = 7.1 Hz, 2H), 4.18–4.09 (m, 2H), 2.61 (s, 3H), 1.29 (t, *J* = 7.1 Hz, 3H). ^13^C NMR (126 MHz, DMSO-*d_6_*) *δ* 164.3, 164.1, 157.7, 151.3, 145.8, 138.9, 136.9, 133.4, 116.7, 113.4, 86.1, 85.3, 60.5, 58.4, 40.3, 25.3, 14.1. ESI-HRMS: calculated for C_18_H_18_ClIN_5_O_5_S_2_ [M+H]^+^, 609.9477; found, 609.9478.

#### 2.2.5. Ethyl 6-(3-(3-((5-[^3^H]chlorothiophen-2-yl-3-*t*)sulfonyl)ureido)azetidin-1-yl)-5-cyano-2- methylnicotinate ([^3^H]AZ12464237)

Precursor **3** (2.5 mg, 4.1 μmol) was dissolved in EtOH (500 μL) with triethylamine (20.0 μL, 144 μmol) and Pd on Ca_2_CO_3_ (2.5 mg, 24 μmol). The sample was placed on a tritium manifold (RC Tritec Ltd., Teufen, Switzerland) and frozen with liquid N_2_. The sample was degassed by way of two freeze/thaw cycles before a partial atmosphere of tritium gas (0.39 mg, 65 μmol, 139 GBq) was introduced. The mixture was allowed to warm to room temperature and was stirred for 90 min. Residual tritium was recovered on a secondary bed and the solvent was removed under a stream of N_2_. Labile tritium was removed by washing with (2 x 0.5 mL) portions of EtOH, which was removed under a stream of N_2_. The crude sample was filtered, washed with EtOH and concentrated under reduced pressure. The residue was redissolved in EtOH (20 mL) to afford a stock solution. The total radioactivity was assessed to be 1558 MBq. LC/MS showed a UV ratio of 37:63 of product to starting material. The crude stock solution was concentrated under reduced pressure, redissolved in DMSO (1 mL) and purified by reversed phase HPLC on a Waters Xbridge C18 5 µm column (100 x 19 mm OBD) using a gradient of MeCN (solvent B) in 0.1% *v*/*v* aqueous trifluoroacetic acid (solvent A); 0 → 2 min (30% B); 2 → 20 min (30% → 95%); 20 → 25 min (95%); Flowrate was 25 mL/min; UV-detection at 292 nm. The collected fractions were merged, concentrated under reduced pressure and redissolved in EtOH (10 mL) to afford a stock solution of [^3^H]AZ12464237 (833 MBq). Product identity was confirmed by analytical HPLC, ^3^H NMR and LC/MS. HPLC method 1 (system 1): Waters Xbridge C18 3.5 μm (100 x 4.6 mm) column with a gradient of MeCN (solvent B) in 10 mM ammonium formate/formic acid pH 3 (solvent A) as eluent. Gradient settings: 0 → 3 min (5% solvent B); 3 → 25 min (5% → 95% solvent B); 25 → 30 min (95% solvent B). Eluent flow rate was 0.6 mL/min and scintillation liquid flow rate was 1.8 mL/min. Retention time of [^3^H]AZ12464237 = 17.20 min; radiochemical purity (RCP) = 96%. HPLC method 2 (system 1): Waters Xbridge C18 3.5 μm (100 x 4.6 mm) column with a gradient of MeCN (solvent B) in 10 mM ammonium bicarbonate buffered with ammonium hydroxide pH 10 (solvent A) as eluent. Gradient settings: 0 → 3 min (5% solvent B); 3 → 25 min (5% → 95% solvent B); 25 → 30 min (95% solvent B). Eluent flow rate was 0.6 mL/min and scintillation liquid flow rate was 1.8 mL/min. Retention time of [^3^H]AZ12464237 = 12.30 min; RCP = 99%. HPLC method 3 (system 2): Platinum C18 100A 5μm (250 × 4.6 mm) column using 45% MeCN (solvent B) in 0.1% trifluoroacetic acid (TFA) in water (solvent A) as eluent. Eluent flow rate was 1 mL/min, and UV detection at 254 nm. Retention time of [^3^H]AZ12464237 = 7.06 min; RCP = 99%. ^3^H NMR (533 MHz, DMSO-*d*_6_) *δ* 7.26 (s). LC/MS (positive ion mode) *m+1/z* (%): 486 (100%), 488 (44.4%), 487 (19.0%), 484 (9.6%).

### 2.3. Cell culture and transfection

HEK 293 (ECACC 8512062) cells were purchased from Sigma-Aldrich (Zwijndrecht, the Netherlands) and cultured in Dulbecco’s modified Eagle’s medium supplemented with 10% fetal bovine serum (FBS), penicillin (100 units/mL), streptomycin (100 μg/mL), and L-glutamine (300 μg/ mL) and maintained at 37 °C, 5% CO2. Stable transfection of HEK 293 cells with human P2Y_12_R (hP2Y_12_R) or rat P2Y_12_R (rP2Y_12_R) was done using lipofectamine 2000 (Invitrogen, San Diego, CA, USA). Plasmid pcDNA3.1(+)-hP2Y_12_R-P2A-eGFP (hP2Y_12_R sequence access number NM_022788.4) and pcDNA3.1(+)-rP2Y_12_R -P2A-eGFP (rP2Y_12_R sequence access number NM_022800.1) were purchased from Genscript (Tokyo, Japan). Briefly, cells were seeded in 24- well plates overnight to reach 70% confluency and were then transfected with the plasmid (0.8 μg/ well) using lipofectamine 2000 according to the manufacturer’s protocol. After 24 h of transfection, cells were transferred to a 6-well plate and kept to recover for 24 h; G-418 (1 mg/mL) (Sigma-Aldrich Zwijndrecht, the Netherlands) was added for positive transfection selection. Media with G- 418 was refreshed every other day for 10 days, and single cells were sorted for eGFP positivity in 96-well plates on a BD FACSARIA Fusion (BD Bioscience, Vianen, The Netherlands). Cell clones were kept to grow and then selected based on the eGFP fluorescence intensity for further expansion. Selected clones were expanded and then analyzed by Fortessa flow cytometry (BD bioscience) for eGFP expression. The HEK 293 cells transfected with the P2X_7_R was prepared according to previously reported protocols.(19)

### 2.4. Real-Time Quantitative Polymerase Chain Reaction (RT-qPCR)

Cells were washed with PBS, after which 250 µL trizol (Life Technologies) was added to dissociate the cells. Chloroform (50 µL) was added to the 250 µL trizol and thoroughly vortexed and then centrifuged for 15 min at 11000 rpm at 4 °C. The water phase (120 µL) was pipetted to a new tube and equal volume of isopropanol 100% (120 µL) was added, thoroughly vortexed, and incubated at -20 °C overnight. Subsequently, samples were centrifuged at 11000 rpm for 10 min at 4 °C. The RNA containing pellet was then washed with 80% ethanol (250 µL), thoroughly vortexed and centrifuged for 9000 rpm for 9 min at 4 °C. The obtained pellet was air dried for 10 minutes and dissolved in DEPC-H_2_O (120 µL). RNA concentration and purity was measured by Nanodrop at 260/280 nm. RNA was then reverse transcribed into DNA using high-capacity cDNA reverse transcription kit-1 (Thermo Fisher scientific). qPCR was run on QuantStudio 3 (Applied Biosystems) using the glyceraldehyde 3-phosphate dehydrogenase (GAPDH) as reference gene and P2Y_12_R as target gene. Samples were run in duplicates with each reaction containing 2 µL or the cDNA, 5 µL SyberGreen mix, 2.3 µL of H_2_O and 0.2 µL of forward/reverse primers. The difference between the Ct of the target gene (P2Y_12_R) and the Ct of the reference gene (GAPDH) ΔCt = Ct _P2Y12R_ − Ct_GAPDH_ is used to obtain the normalized amount of target (Nt), which corresponds to 2−ΔCt. The Nt reflects the relative amount of target transcripts with respect to the expression of the GAPDH. Table 1 shows the used primer sequences.

**Table 1.**
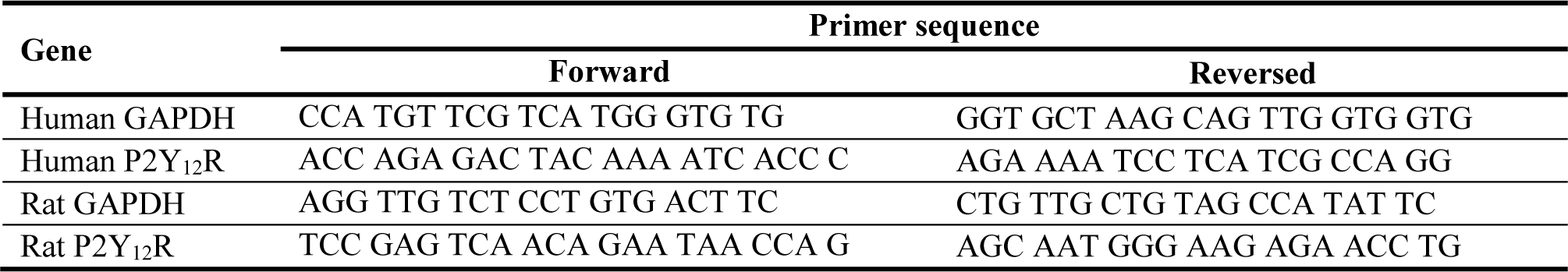
Forward and reverse primer sequences used in qPCR assays.

### 2.5. Membrane preparation

For membrane preparation, cells were washed twice with PBS and collected using a scraper and centrifuged for 10 min at 1720 rpm at 4 °C. The cells were resuspended in ice-cold lysis buffer (50 mM Tris/HCl, 140 mM NaCl, 2 mM EDTA, cOmplete™ ULTRA Tablets, Mini, EASY pack Protease Inhibitor Cocktail 1 tablet/10 mL buffer, pH 7.4) and homogenized (while kept on ice) using a Branson Sonifier 250 (2 x 15 sec at full speed). The homogenates were centrifuged in an Optima XE-90 Ultracentrifuge (Beckman Coulter Inc., Indianapolis, IN, USA) at 20000 rpm for 30 min at 4 °C. The supernatant was discarded and the pellets were resuspended in lysis buffer and centrifuged a second time with the same settings. The supernatant was discarded and the pellets were resuspended in ice-cold Tris/HCl buffer (50 mM Tris/HCl, 140 mM NaCl, 2 mM EDTA, pH 7.4) The protein concentration was determined with a BCA protein kit, and bovine serum albumin as standard (ThermoFisher Scientific, Breda, the Netherlands). Afterward, the membranes were aliquoted at a concentration of 1μg/μL and stored at -80 °C until they were used in binding experiments.

### 2.6. Binding assays

#### 2.6.1. Radioligand binding - general conditions

All binding assays were carried out by incubation in Na/K-phosphate buffer (50 mM Na_2_HPO_4_/KH_2_PO_4_, 100 mM NaCl, 5 mM MgCl_2_, 0.1% Tween-20, pH 7.4) at room temperature with agitation (200 rpm). This buffer was used for all assays except in the buffer screening. The total volume per binding tube was 500 µL. Addition order was as follows; 100 µL solution of competitor (tubes for non-specific binding (NSB) assessment) or only buffer (total binding tubes), 100 µL of radioligand solution and 300 µL of membrane suspension. For all assays the incubations were terminated by vacuum filtration onto GF/B glass-fiber filters (Alpha Biotech Ltd., Killearn, United Kingdom) pre-soaked in 5% aqueous solution of PEI (soaked for 1 h prior to use) using a Brandel cell harvester (Brandel, Gaithersburg, MD, USA), followed by three rapid washes (5 mL) with ice-cold Na/K-phosphate buffer (50 mM, pH 7.4), if not stated otherwise. The filters were dried, cut and collected to vials, to which 5 mL scintillation liquid (PerkinElmer, Drachten, The Netherlands) was added. Radioactivity bound to the filter was counted in an Hidex 300 SL automatic liquid scintillation counter (Hidex, Turku, Finland) for 10 min per sample. Each assay was performed in three different experiments and each data point has been carried out in triplicate.

#### 2.6.2. Screening of binding buffer

Membranes of hP2Y_12_R-HEK (10 µg) were incubated for 1 h with [^3^H]AZ12464237 (5.5 nM) using the conditions described above in different binding buffers based on either Tris/HCl (50 mM Tris, 100 mM NaCl, 5 mM MgCl_2_, pH 7.4), HEPES (50 mM HEPES, 100 mM NaCl, 5 mM MgCl_2_, pH 7.4) or Na/K-phosphate (50 mM, 100 mM NaCl, 5 mM MgCl_2_, pH 7.4) buffer with different additives (BSA, Tween-20, Tween-80 or Triton X-100). NSB was determined by co-incubation with either elinogrel (20 µM) or ticagrelor (10 µM). The washing buffer was the same as the basis of the binding buffer *i.e.*, Tris/HCl (50 mM Tris, pH 7.4), HEPES (50 mM HEPES, pH 7.4) or Na/K-phosphate (50 mM Na_2_HPO_4_/KH_2_PO_4_, pH 7.4) buffer, respectively without any additional additives.

#### 2.6.3. Kinetic measurements of [^3^H]AZ12464237 binding to hP2Y_12_R

Association studies were performed by incubating different concentrations of [^3^H]AZ12464237 (0.5, 1.5, 5.0 and 15 nM) with hP2Y_12_R-HEK (20 µg). The studies were started at varying time points by adding the membranes and thereafter harvested at the same time. NSB was determined by co-incubation with elinogrel (20 µM).

Dissociation studies were performed by allowing [^3^H]AZ12464237 (1.5 nM) and hP2Y_12_R-HEK (20 µg) to reach equilibrium *i.e.*, 1 h. Afterward, elinogrel (20 µM) was added at different time points to compete with [^3^H]AZ12464237 for binding to the receptor.

#### 2.6.4. Saturation binding of [^3^H]AZ12464237 to hP2Y_12_R and rP2Y_12_R

A range of different concentrations of [^3^H]AZ12464237 (0.5, 1.0, 2.0, 4.0, 6.0, 8.0, 10, 12, 20, 40, 60 and 90 nM) were incubated with hP2Y_12_R-HEK membranes (20 µg) or rP2Y_12_R-HEK membranes (10 µg) for 1 h. NSB was determined by co-incubation with elinogrel (20 µM). The TB never exceeded 10% of the added concentrations of [^3^H]AZ12464237. Incubations were terminated as described above.

#### 2.6.5. Competitive displacement of [^3^H]AZ12464237 binding to hP2Y_12_R and rP2Y_12_R

A series of concentrations (10 pM–100 μM) of different ligands were incubated with a single concentration of [^3^H]AZ12464237 (5.0 nM) and hP2Y_12_R-HEK membranes (20 µg) or [^3^H]AZ12464237 (15 nM) and rP2Y_12_R-HEK membranes (10 µg) for 1 h. Incubations were terminated as described above.

### 2.7. Data Analysis

All binding assay data were analyzed using GraphPad Prism 8.0.2 (San Diego, CA, USA). Experiments were carried out in triplicates in three independent studies and all values were averaged to obtain mean ± SEM if not otherwise stated. *p*-values of less than 0.05 were considered statistically significant. The data were fit and analyzed using the following equations:

Association studies with global fitting using four different concentrations of the radioligand:

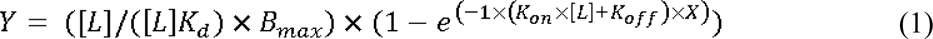

Where [*L*] is the concentration of radioligand in nM, *K_d_* is the equilibrium dissociation constant, *B_max_* is the maximum binding at equilibrium at maximal radioligand concentration in the units of *Y*, *K_on_* is the association rate constant in M^-1^ min^-1^ and *K_off_* is the dissociation rate constant in min^-1^.

Association data for each radioligand concentration were also fitted to an exponential one phase association to calculate the observed rate constant *k_obs_*:

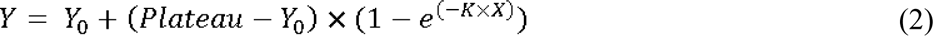

Where *Y_0_* is the value of *Y* when *X* (time) is 0, the plateau is the value of *Y* at infinite times, *K* is the observed rate constant (*k_obs_*) for the tested radioligand concentration. Next, the *k_obs_* values were plotted against the radioligand concentrations. The slope of this plot corresponds to *K_on_* and the *Y*- intercept equals to *K_off_*. Afterward, *K_d_* can be estimated from equation 3.

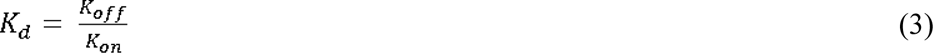

Dissociation data were fitted and analyzed with one phase exponential decay:

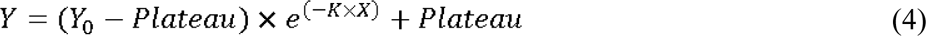

Where K is the dissociation rate constant (*K_off_*).

Data from saturation binding experiments were fitted to equation 5.

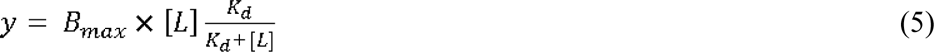

Where [L] represents increasing concentrations of the radioligand.

Competition binding experiments were analyzed by curve fitting according to equation 6.

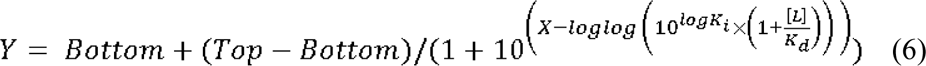

Where *Top* and *Bottom* are plateaus in the units of *Y*, [*L*] is the concentration of the radioligand in nM and *K_d_* is the affinity value of the radioligand in nM.

### 2.8. Autoradiography and immunohistochemistry

#### 2.8.1. Tissues and Animals

For the in vitro autoradiography studies 10um thick cryosections of frozen cell pellets or mouse brains were used. The cell pellets were obtained from Chinese hamster ovary (CHO) cells overexpressing (oe) either P2Y_12_R or a control protein which were obtained from Rochès cell-line repository. Cryosections from 4-month old female Wild-Type (WT) and P2Y_12_R knock-out (KO) mice were kindly provided by Dr. Denes (Institute of Experimental Medicine, Budapest, Hungary) and the line has been previously studied and characterized.(27) All sections were generated with a Leica CM3050 cryostat (Leica Biosystems, Nussloch, Germany) at approximately -15°C chamber and -17 °C object temperature and were then mounted on Histobond+ microscope slides (Marienfeld Laboratory Glassware, Lauda-Königshofen, Germany). The sections were then air-dried for maximum 3 h at room temperature prior storage at -20 °C until used.

#### 2.8.2. *In vitro* autoradiography

Cryosections were incubated in assay buffer (20 mM Tris-HCl buffer, pH 7.4) containing [^3^H]AZ12464237 (10 nM) for 30 min. The NSB was assessed by co-incubation with AZ12464237, ticagrelor, cangrelor and a structurally unrelated compound for the CHO sections, and AZ12464237, ticagrelor, cangrelor, staurosporine, rosiglitazone and an unrelated compound for the mouse tissue sections. All the compounds were used at 10 μΜ concentration. After the incubation, all sections were rinsed (3 x 1.5 min) in a cold assay buffer and dipped once in cold distilled water. The sections were dried for at least 3 h and thereafter exposed to tritium sensitive film (TR2025, Fujifilm, Minato City, Japan) shielded in exposure cassettes (RPA-510, GE Healthcare, Chicago, IL, USA) along with a ^3^H-Microscale (American Radiolabeled Chemicals Inc., St. Louis, MO, USA) for five days. The imaging plate was scanned with 25-μm resolution in a high-resolution plate scanner (BAS-5000, Fujifilm, Minato City, Japan) and the autoradiography was analyzed using MCID^TM^ 7.0 Software (Imaging Research Inc., St. Catharines, ON, Canada). The total amount of radioligand bound to the brain areas of interest was expressed as fmol of bound radioligand per mg of protein.

### 2.9. Immunohistochemistry

Sections were fixed for 15 min in 4% paraformaldehyde (PFA) and rinsed with PBS. The sections were blocked for 40 min in the blocking solution containing 1% of bovine serum albumin (BSA), ovalbumin and goat serum. The CHO cell pellet sections were incubated with the primary antibodies for 2 h at room temperature, whereas the mouse brain sections overnight at 4 °C, along with 0.25% Triton. Upon rinsing with PBS the slices were incubated for 1 h at room temperature with the secondary antibody mix in the blocking solution. Lastly, the sections were counterstained with DAPI, which was followed by incubation with 0.3% Sudan Black, only for the case of the mouse brain sections for the reduction of autofluorescence, and they were mounted with Aqua-Poly/Mount (Polysciences, 18606-20) and cover slipped. Large scanned images of the CHO and mouse sections were acquired with the VS200 Slide Scanner (OlympusLife Science, Waltham, MA, USA). Primary antibodies: 1:300 P2Y_12_R (Abcam, 300140), 1:500 Iba1 (Synaptic systems, 234009). Secondary antibodies: 1:200 goat-anti-rabbit-488 (Invitrogen, A11034), 1:200 donkey-anti-chicken-647 (Jackson ImmunoResearch,703-605-155).

## 3. Results

### 3.1. Chemistry

The radioligand was envisioned to be obtained via Pd-mediated tritiation of a corresponding iodine precursor. Thus, compound **3** was synthesized (Scheme 1), and in order to confirm the success of the radiolabeling, the non-radioactive analog of AZ12464237 was synthesized as reference according to previously described procedures. The ^3^H-labeled product was characterized by HPLC (Figure 2), LC/MS and NMR. NMR experiments verified that ^3^H-labeling occurred in the 4- position of the thiophene of **3** to a high extent. A precise level of incorporation could not be determined in this case due to the lack of a clear residual proton peak in the 1D ^1^H NMR spectrum of [^3^H]AZ12464237. However, the 1D ^3^H NMR spectrum shows only one peak, suggesting that ^3^H was only introduced in one position of the molecule. In the end, [^3^H]AZ12464237 was afforded in an activity yield of 833 MBq, a molar activity (A*_m_*) of 969 kBq/nmol, and a high radiochemical purity (RCP).

**Scheme 1.**
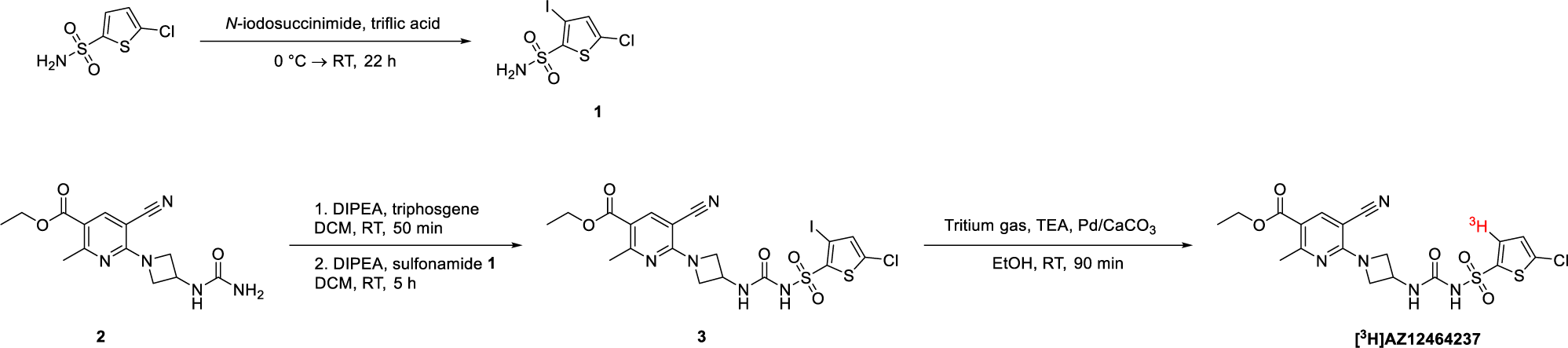
Synthesis of precursor and subsequent radiolabeling to afford [^3^H]AZ12464237. DCM, dichloromethane; DIPEA, *N*,*N*- diisopropylethylamine; EtOH, ethanol; TEA, triethylamine; RT, room temperature.

**Figure 2.**
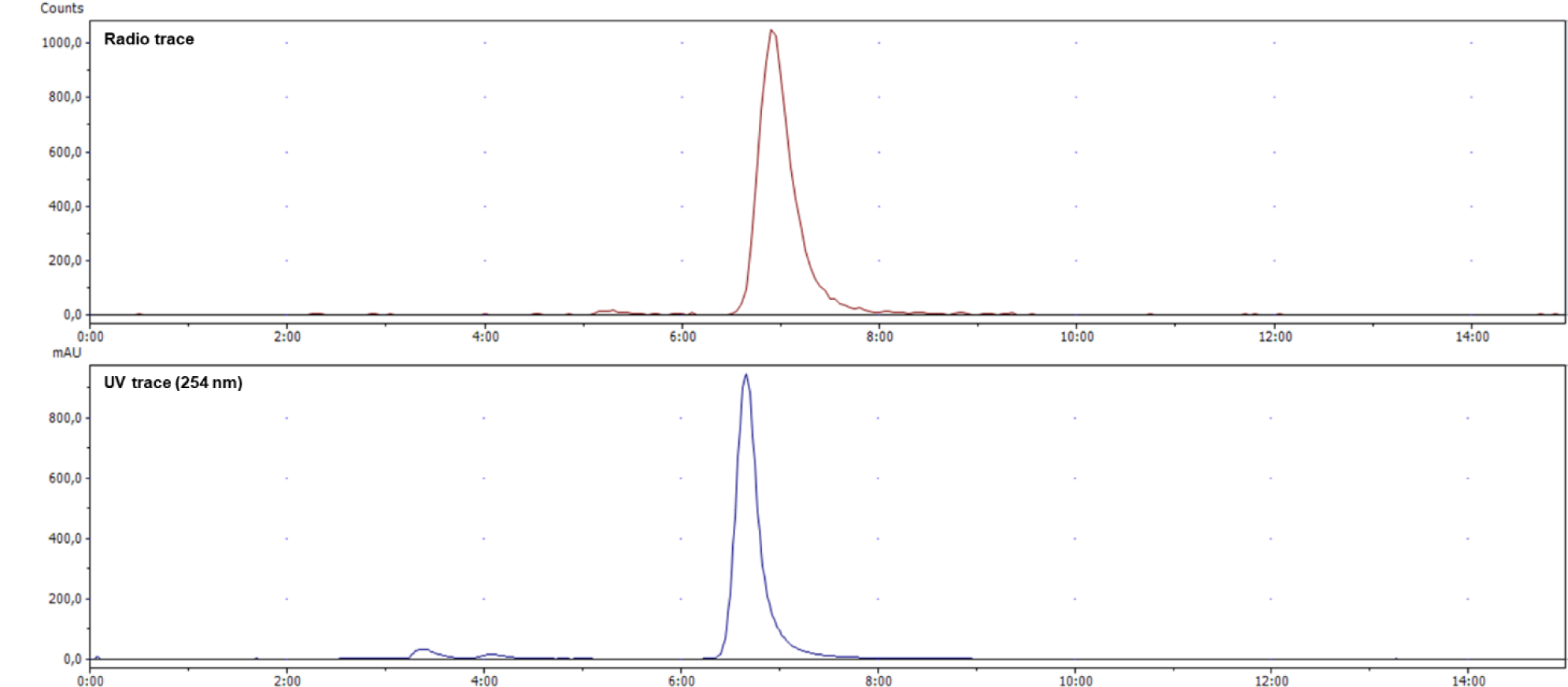
Analytical HPLC chromatogram of [^3^H]AZ12464237, co-injected with unlabeled reference compound (AZ12464237). HPLC conditions: Platinum C18 100A 5μm (250 × 4.6 mm) column using MeCN/H_2_O/trifluoroacetic acid (TFA) (45:55:0.1, *v/v/v*) as eluent with UV detection at 254 nm. Eluent flow rate was 1 mL/min and scintillation liquid flow rate was 0.3 mL/min. Retention time = 7.06 min; RCP = 99%.

### 3.2. Development of *in vitro* binding assay on membranes preparations

HEK-293 cells were successfully stably transfected with the hP2Y_12_R. The transfection efficiency was confirmed by qPCR and flow cytometry to detect eGFP expression (Figure 3A and B). Membrane preparations from the transfected cells were used for the assay development and pharmacological characterization of the binding of [^3^H]AZ12464237 to hP2Y_12_R. We started off by performing a preliminary screening of different buffers to find a suitable binding buffer to detect [^3^H]AZ12464237 binding to hP2Y_12_R. The investigated buffers were based on either Tris-HCl (50 mM), HEPES (50 mM) or Na/K-phosphate (50 mM), with NaCl (100 mM), MgCl_2_ (5 mM) and different additives *i.e.*, bovine serum albumin (BSA), Tween-20, Tween-80 and Triton X-100 (Figure 3C–E). The Na/K-phosphate buffer with 0.1% Tween-20 (Figure 3E) turned out to be the most promising buffer to use in binding studies, with a sufficient number of counts for total binding (TB) and acceptable levels of non-specific binding (NSB, 16% of the TB). This buffer was used in all of the following experiments involving binding on membrane preparations.

**Figure 3.**
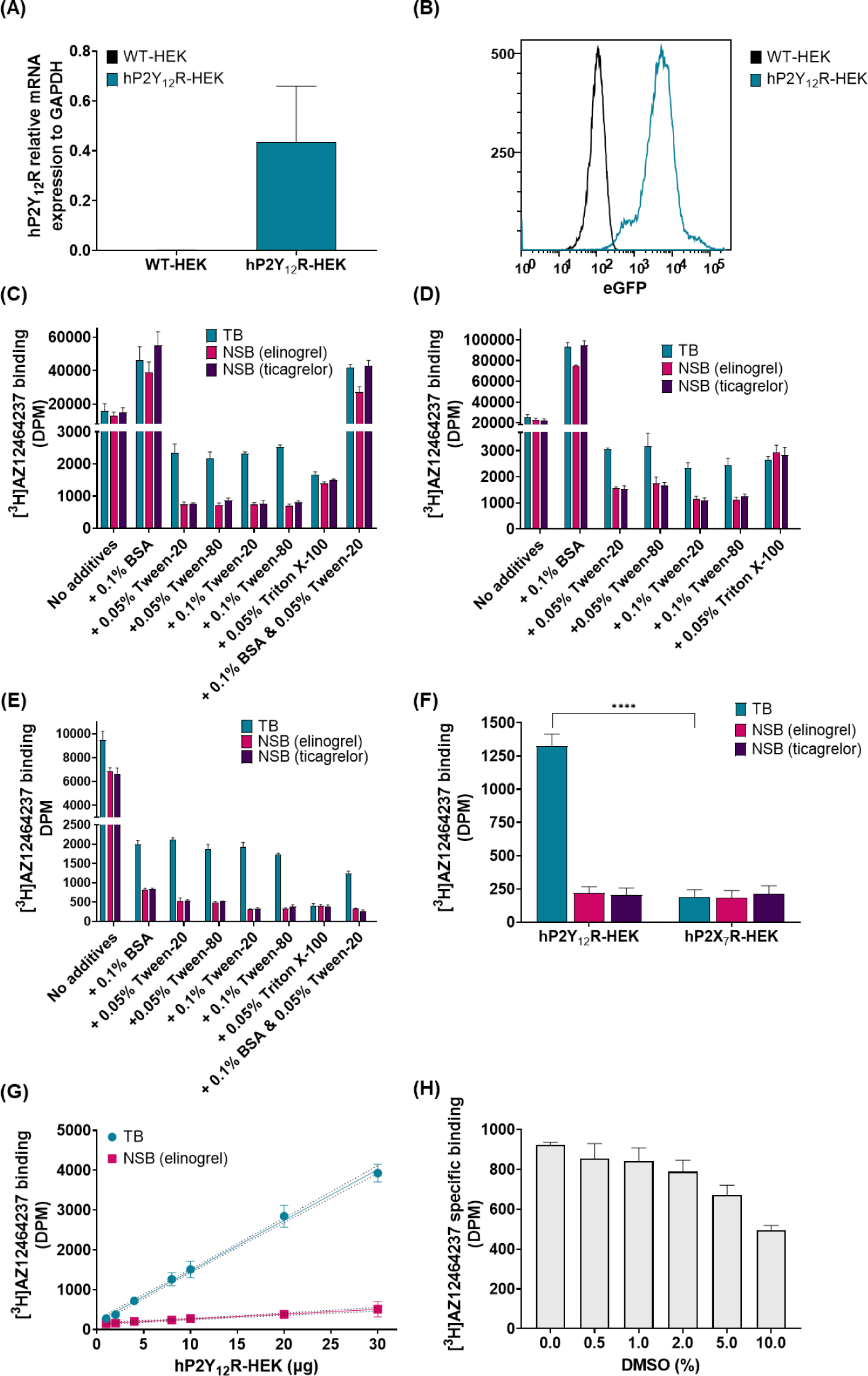
Transfection efficiency of hP2Y_12_R-HEK 293 cells and optimization of assay conditions for [^3^H]AZ12464237 binding to membrane preparations of the same cells. **(A)** Relative qPCR analysis of the mRNA expression of hP2Y_12_R in WT (black bar) and transfected (blue-green bar) HEK 293 cells. **(B)** The eGFP expression of WT (black line) and transfected (blue-green line) HEK 293 cells as detected by flow cytometry. **(C)** Screening of binding buffers with different additives (as indicated in graph). Buffer was based on Na/K-phosphate (50 mM Na_2_HPO_4_/KH_2_PO_4_, 100 mM NaCl, 5 mM MgCl_2_, pH 7.4). BSA = Bovine serum albumin. **(D)** Screening of binding buffers with different additives (as indicated in graph). Buffer was based on Tris-HCl (50 mM Tris, 100 mM NaCl, 5 mM MgCl_2_, pH 7.4). **(E)** Screening of binding buffers with different additives (as indicated in graph). Buffer based on HEPES (50 mM HEPES, 100 mM NaCl, 5 mM MgCl_2_, pH 7.4) **(F)** Control experiments to determine the specificity of binding to hP2Y_12_R. Membranes prepared from HEK 293 cells transfected with the hP2X_7_R were used as a negative control. Blue-green bars represents total binding (TB), whereas pink and dark purple bars represent assessment of NSB by co-incubation with elinogrel (20 μM) and ticagrelor (10 μM), respectively. The binding buffer was Na/K-phosphate-based with 0.1% Tween-20. **(G)** Binding of [^3^H]AZ12464237 with increasing amount of hP2Y_12_-HEK membranes (0.5–30 μg per 500 uL total volume in binding tube). Blue-green line represents TB and the pink line represents NSB determined with elinogrel (20 μM). The binding buffer was Na/K-phosphate-based with 0.1% Tween-20. **(H)** The influence of DMSO on binding of radioligand. Incubation of [^3^H]AZ12464237 and hP2Y_12_R-HEK for 1 h at room temperature in the presence of different concentrations of DMSO (0.002%, 0.5%, 1%, 2%, 5% and 10%) of the total volume in the binding tube. The binding buffer was Na/K-phosphate-based with 0.1% Tween-20. NSB was assessed by co-incubation with elinogrel (20 μM). Data is presented as mean ± SEM form three different experiments and each data point has been carried out in triplicate. Significant difference was assessed by Student’s *t*-test (\*\*\*\**p*-value <0.0001).

As control, the binding of [^3^H]AZ12464237 to hP2Y_12_R-HEK membranes was compared to its binding to a membrane preparation from HEK-293 cells transfected with the human ion-channel P2X_7_ (hP2X_7_R). As shown in Figure 3F, a 7-fold higher binding of [^3^H]AZ12464237 was detected with hP2Y_12_R-HEK membranes. Also, the binding of [^3^H]AZ12464237 when using the membranes with high hP2X_7_R expression equaled the levels of NSB with the hP2Y_12_R-HEK membranes. Moreover, the specific binding (SB) of [^3^H]AZ12464237 to hP2Y_12_R was proportional to the amount of membrane used (Figure 3G). At all tested protein concentrations, the percentage TB compared to the total amount of added radioligand was below 10%, hence avoiding ligand depletion. The highest levels of bound radioligand was 4%, and was observed at the highest protein concentration studied (30 μg/tube; 60 μg/mL). For subsequent binding studies, a protein amount of 20 μg (40 μg/mL) was selected. This amount provided an appropriate number of counts and ensured a reliable window between TB and NSB for the pharmacological characterization studies.

As a final experiment before more detailed pharmacological characterization, the influence of DMSO on the binding of [^3^H]AZ12464237 to hP2Y_12_R was studied (Figure 3H). DMSO concentrations of up to 2% of the total volume had only a minor effect on the binding of [^3^H]AZ12464237. At the 2% DMSO concentration the SB was 85% of the SB obtained with the lowest possible DMSO content (0.02% from the stock of the radioligand). In light of this, the level of DMSO was kept below 2% in the subsequent competitive binding studies.

### 3.3. *In vitro* characterization of [^3^H]AZ12464237 binding to hP2Y_12_R

After establishing appropriate assay conditions, the binding of [^3^H]AZ12464237 to hP2Y_12_R was fully characterized in kinetic binding studies to study its association and dissociation, as well as in saturation binding experiments. Association studies were performed using four different concentrations of the radioligand *i.e.*, 0.5, 1.5, 5.0 and 15 nM. This concentration range was based on a rough estimation of the expected *K_d_* (2–3 nM). The binding of [^3^H]AZ12464237 to hP2Y_12_R was terminated at designated time points between 2 and 180 min. As for previous studies, elinogrel was used for the assessment of NSB. The level of binding remained constant during the course of 3h at room temperature for all concentrations tested. The obtained association data for the first 30 min were fitted in two ways. Firstly, global fitting using equation 1 resulted in a *K_on_* of 0.0698 ± 0.0450 nM^-1^ min^-1^ and a *K_off_* = 0.304 ± 0.024 min^-1^. The kinetic *K_d_* derived from these parameters is 4.35 ± 0.21 nM. Secondly, the association data for each concentration was fitted to an exponential one phase association. In this way *k_obs_* was obtained, which in turn can be plotted against the concentration of radioligand (Figure 4B). This approach resulted in the following values; *K_on_* = 0.045 ± 0.006 nM^-1^ min^-1^, *K_off_* = 0.419 ± 0.045 min^-1^, *K_d_* = 9.35 nM.

**Figure 4.**
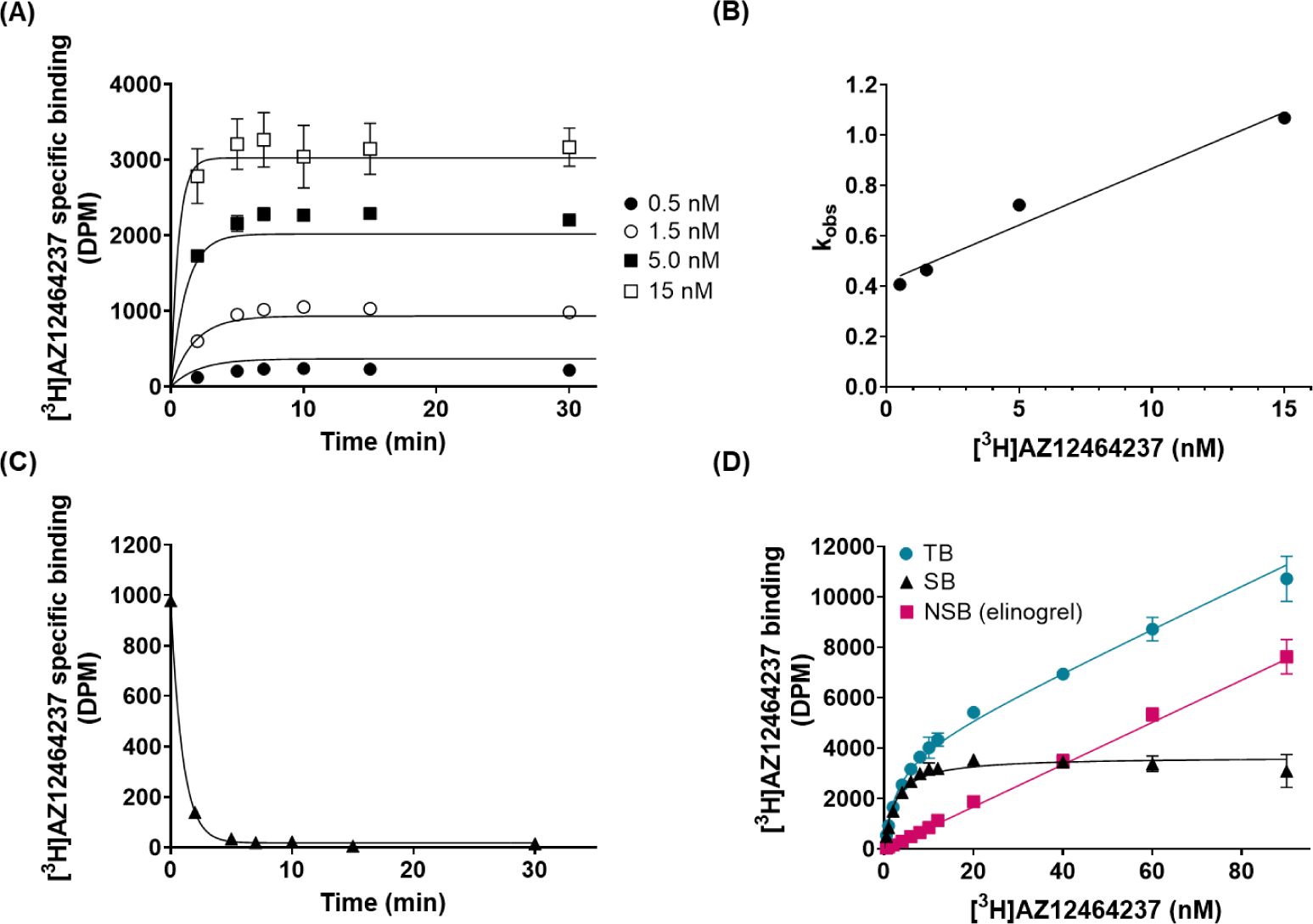
Characterization of [^3^H]AZ12464237 binding to membrane preparations of HEK 293 cells transfected with the hP2Y_12_R. (**A**) Association studies performed with four different concentrations of [^3^H]AZ12464237; 0.5 nM (filled circles), 1.5 nM (open circles), 5.0 nM (filled squares) and 15 nM (open squares). (**B**) The *k_obs_* values obtained from association data fitted to exponential one phase association plotted against the concentration of [^3^H]AZ12464237. (**C**) Dissociation studies were performed by incubating [^3^H]AZ12464237 (1.5 nM) with hP2Y_12_R-HEK for 1 h to reach equilibrium before dissociation by addition of elinogrel (20 μM) at different time points. (**D**) Saturation binding studies with increasing concentrations of [^3^H]AZ12464237 (0.50–90 nM) in the presence or absence of elinogrel (20 μM) to assess NSB. The blue-green line represents TB, the black line corresponds to the specific binding (SB) and the pink line is the NSB. Graphs are representatives of three experiments and each data point has been carried out in triplicates.

Next, dissociation studies were carried out at one concentration of radioligand. Here, [^3^H]AZ12464237 was incubated with hP2Y_12_R-HEK membranes for 1h at room temperature to reach equilibrium, followed by addition of an excess of elinogrel (20 μM) at different time points between (2–30 min). The data was fitted by one phase exponential decay (equation 4), resulting in a *K_off_* = 0.928 ± 0.168 min^-1^. The associated target residence time, which is defined as 1/*K_off_,* results in a value of 1.10 ± 0.22 min. Using the obtained *K_off_* combined with the *K_on_* from association studies (global fitting) results in a *K_d_* = 13.3 ± 2.40 nM.

Once the kinetic parameters of [^3^H]AZ12464237 had been determined, saturation binding studies were carried out (Figure 4D). The radioligand, in increasing concentrations (0.5–90 nM), was incubated with the membranes for 1 h. The data was fitted to equation 5 and revealed a *K_d_* of 3.12 ± 0.70 nM and a maximal binding number of binding sites (*B_max_*) of 3.46 ± 0.55 pmol/mg. Table 2 summarizes the binding parameters obtained in the different studies; the *K_d_* values of [^3^H]AZ12464237 binding to hP2Y_12_R-HEK membranes from different experiments range from 3.12 – 13.3 nM, clearly showing high affinity binding of the radioligand. At a concentration around the *K_d_* value, [^3^H]AZ12464237 demonstrated a SB of ∼90% (Figure 3D).

**Table 2.** Binding parameters obtained from kinetic studies and saturation binding studies.

| Type of assay | $K_{on}$<br>(nM<sup>-1</sup> min<sup>-1</sup>) | $K_{off}$<br>(min<sup>-1</sup>) | $K_d$<br>(nM) |
| --- | --- | --- | --- |
| Association GF | 0.0698 ± 0.045 | 0.304 ± 0.024 | 4.35 ± 0.21 |
| Association OP | 0.045 ± 0.006 | 0.419 ± 0.045 | 9.35<sup>a</sup> |
| Dissociation | - | 0.928 ± 0.168 | 13.3 ± 2.40 |
| Saturation | - | - | 3.12 ± 0.70 |

Notes: <sup>a</sup> $K_d$ was derived from the mean values of $K_{on}$ and $K_{off}$ obtained from the association studies analyzed by exponential one phase association. GF = association kinetics analyzed by global fitting; OP = association kinetics analyzed by exponential one phase association.

### 3.4. Displacement of [^3^H]AZ12464237 binding with reference ligands for P2Y receptors

To pharmacologically characterize the binding of [^3^H]AZ12464237 to hP2Y_12_R-HEK membranes, [^3^H]AZ12464237 was used in competitive displacement studies against a group of commercially available P2Y receptor ligands (Figure 5; Table 3). Five of these ligands are reported P2Y_12_R ligands, including one agonist (2MeSADP) and four antagonists (AZD1283, elinogrel, PSB0739 and ticagrelor). AZD1283 and elinogrel, which share the highest structural similarity with [^3^H]AZ12464237, also showed the highest *K_i_*-values; 2.66 ± 0.60 nM (AZD1283) and 7.00 ± 0.54 nM (elinogrel). The specificity of binding was further investigated by testing a few P2Y antagonists designed to target other P2Y subtypes, such as P2Y_6_R, P2Y_13_R and P2Y_14_R. As expected, these ligands did not show any displacement or very low affinity *e.g.*, MRS2211 with a *K_i_* of 3.24 ± 1.85 µM. Figure 5 shows representative binding curves for four of the studied compounds; elinogrel, ticagrelor, MRS2211 (P2Y_13_R antagonist) and PPTN·HCl (P2Y_14_R antagonist).

**Figure 5.**
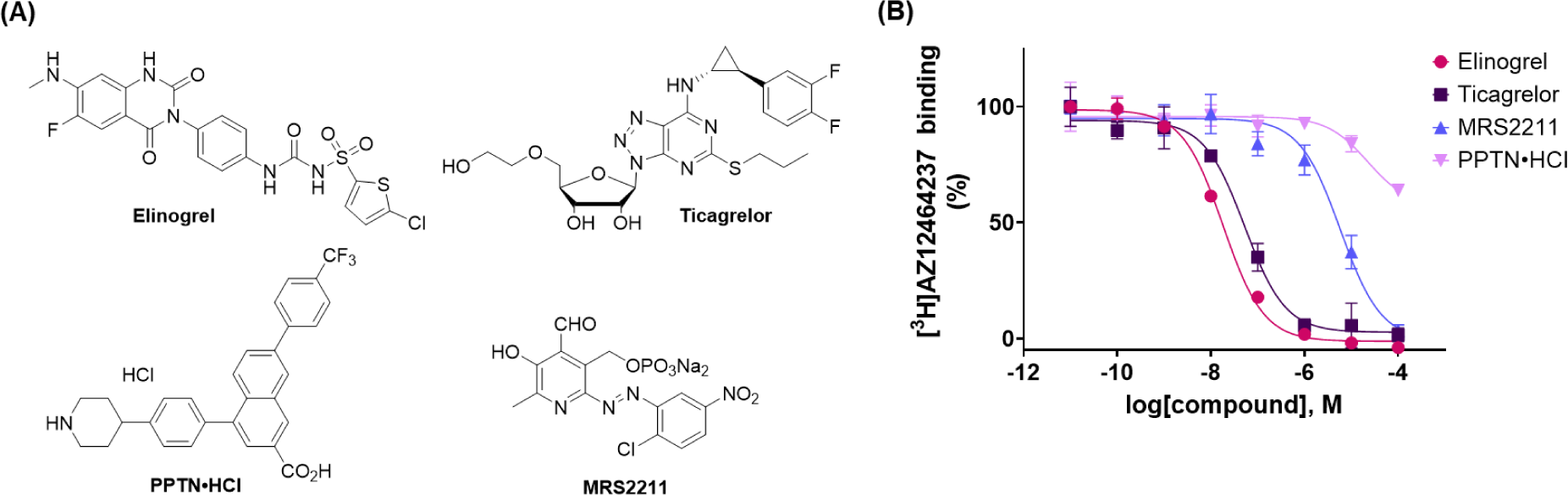
Displacement of [^3^H]AZ12464237 binding to hP2Y_12_R-HEK using different P2Y ligands as competitors. (**A**) Chemical structures of elinogrel (P2Y_12_R antagonist), ticagrelor (P2Y_12_R antagonist), MRS2211 (P2Y_13_R antagonist) and PPTN·HCl (P2Y_14_R antagonist). (**B**) Representative curves obtained from competitive binding of [^3^H]AZ12464237 and elinogrel (pink), ticagrelor (dark purple), MRS2211 (light blue) and PPTN·HCl (light pink) in concentrations ranging from 10 pM to 100 μM. Each experiment has been carried out in triplicates. Graphs are representatives of three experiments and each data point has been carried out in triplicates.

**Table 3.**
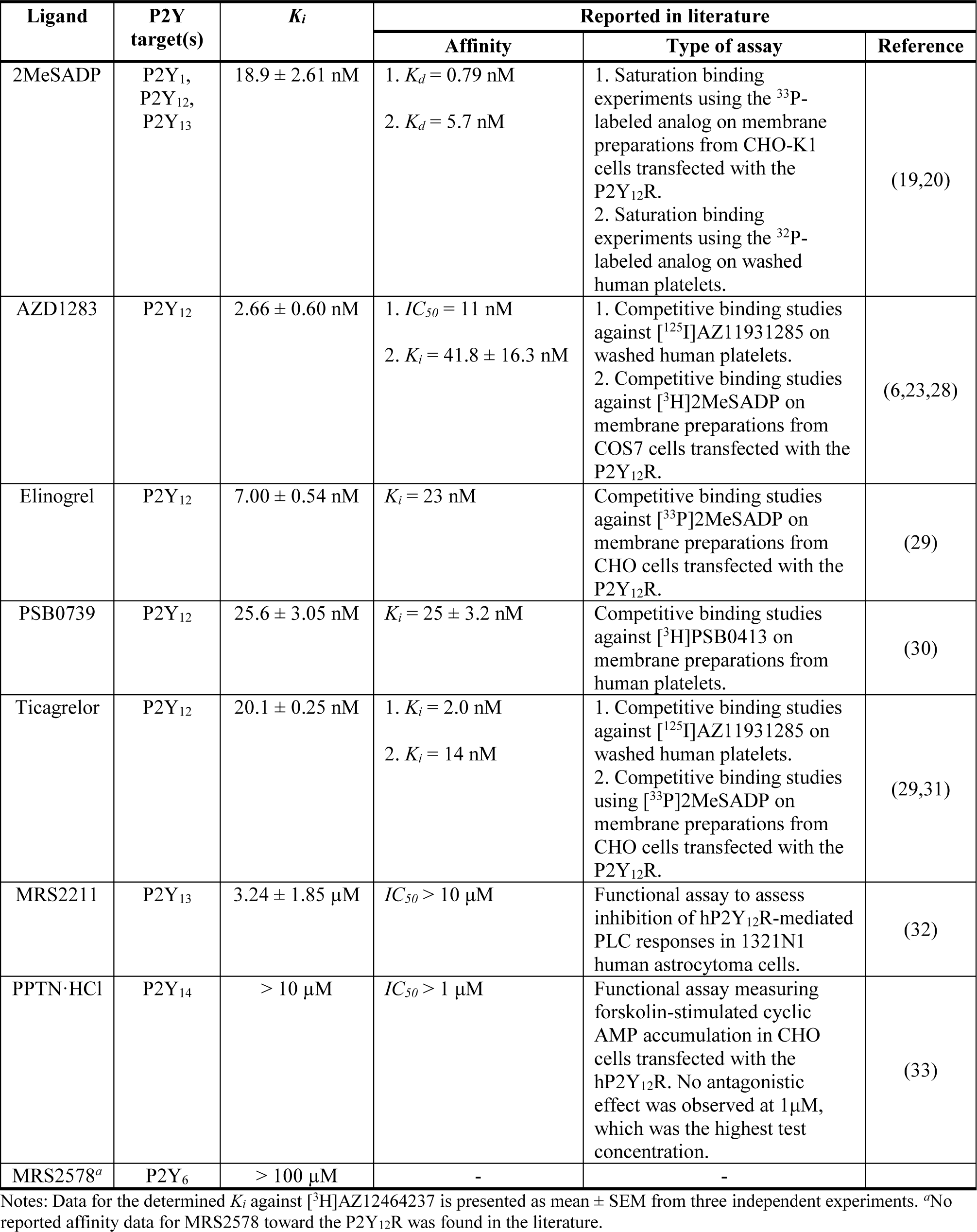
Determined *K_i_*-values for the P2Y_12_R for a set of reported P2Y ligands compared to reported affinity in literature.

### 3.5. Characterization of [^3^H]AZ12464237 binding to rP2Y_12_R

To assess any potential species differences, the binding of the radioligand toward the rat P2Y_12_R (rP2Y_12_R) was also explored. As for the hP2Y_12_R, membranes were prepared from HEK 293 cells, stably transfected with rP2Y_12_R as determined by qPCR (Figure 6A). These membrane preparations were incubated in saturation binding experiments (Figure 6B) with a range of increasing radioligand concentrations (0.50–150 nM). The total binding never exceeded 10% of the added concentrations of [^3^H]AZ12464237. The studies revealed a *K_d_* of 16.6 ± 3.37 nM and a *B_max_* of 74.7 ± 13.2 pmol/mg for the binding of [^3^H]AZ12464237 to rP2Y_12_R. Competitive [^3^H]AZ12464237 binding displacement studies (Figure 6C) were also carried out and revealed that both elinogrel (*K_i_* = 55.03 ± 14.3 nM) and ticagrelor (*K_i_* = 48.8 ± 15.2 nM) show comparable affinities for rP2Y_12_R and hP2Y_12_R.

**Figure 6.**
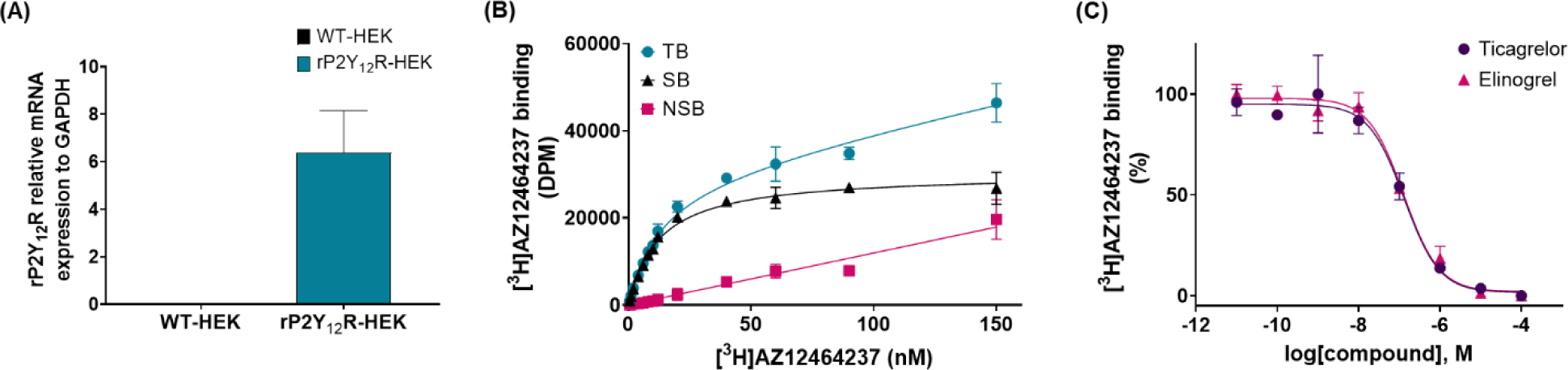
Binding studies of [^3^H]AZ12464237 and competitors to membrane preparations of HEK 293 cells transfected with the rP2Y_12_R. (**A**) Transfection of HEK 293 cells with the rP2Y_12_R. Relative qPCR analysis of the mRNA expression of rP2Y_12_R in WT (black bar) and transfected (blue-green bar) HEK 293 cells. (**B**) Saturation binding studies with increasing concentrations of [^3^H]AZ12464237 (0.50–150 nM) in the presence or absence of elinogrel (20 μM) to assess NSB. The blue-green line represents TB, the black line corresponds to the SB and the pink line is the NSB. (**C**) Competitive binding of [^3^H]AZ12464237 with ticagrelor (dark purple) and elinogrel (pink), respectively in concentrations ranging from 10 pM to 100 μM. Graphs are representatives of three experiments and each data point has been carried out in triplicates.

### 3.6. *In vitro* autoradiography on CHO cell pellets and mouse brain tissue

As a first proof of concept, in order to evaluate the suitability of the radioligand as a tool for autoradiography, the binding was studied on sections derived from frozen CHO cell pellets overexpressing either the hP2Y_12_R or a control protein, obtained from Rochès cell-line repository. The expression of hP2Y_12_R in sections of a pellet of P2Y_12_R-expressing CHO cells was confirmed with immunolabeling against P2Y_12_R, as shown in Figure 7A (top row), whereas the presence of the protein was not observed on the sections derived from the control CHO cell pellets (Figure 7A, bottom row). This differential expression of P2Y_12_R, should allow the evaluation of the tracer’s ability to bind P2Y_12_R. The sections were then incubated with the radioligand only, to assess the TB, or along with different blockers to assess the NSB (representative images, Figure 7B). As shown in Figure 7C, on sections from overexpressing hP2Y_12_R CHO cell pellets, [^3^H]AZ12464237 binding was reduced significantly upon co-incubation with AZ12464237 (88% decrease), ticagrelor (86% decrease) and cangrelor (71% decrease). Whereas, co-incubation with an unrelated compound, with no reported antagonistic effect toward the receptor, did only result in a 11% decrease of binding. On the contrary, when sections from the control CHO cell pellets were incubated with [^3^H]AZ12464237, the binding was 82% lower compared to that quantified on the hP2Y_12_R CHO cell pellet sections. These results, along with the pharmacological inhibition of the binding to the sections of the pellet of hP2Y_12_R CHO cells point toward the binding of [^3^H]AZ12464237 to hP2Y_12_R.

**Figure 7.**
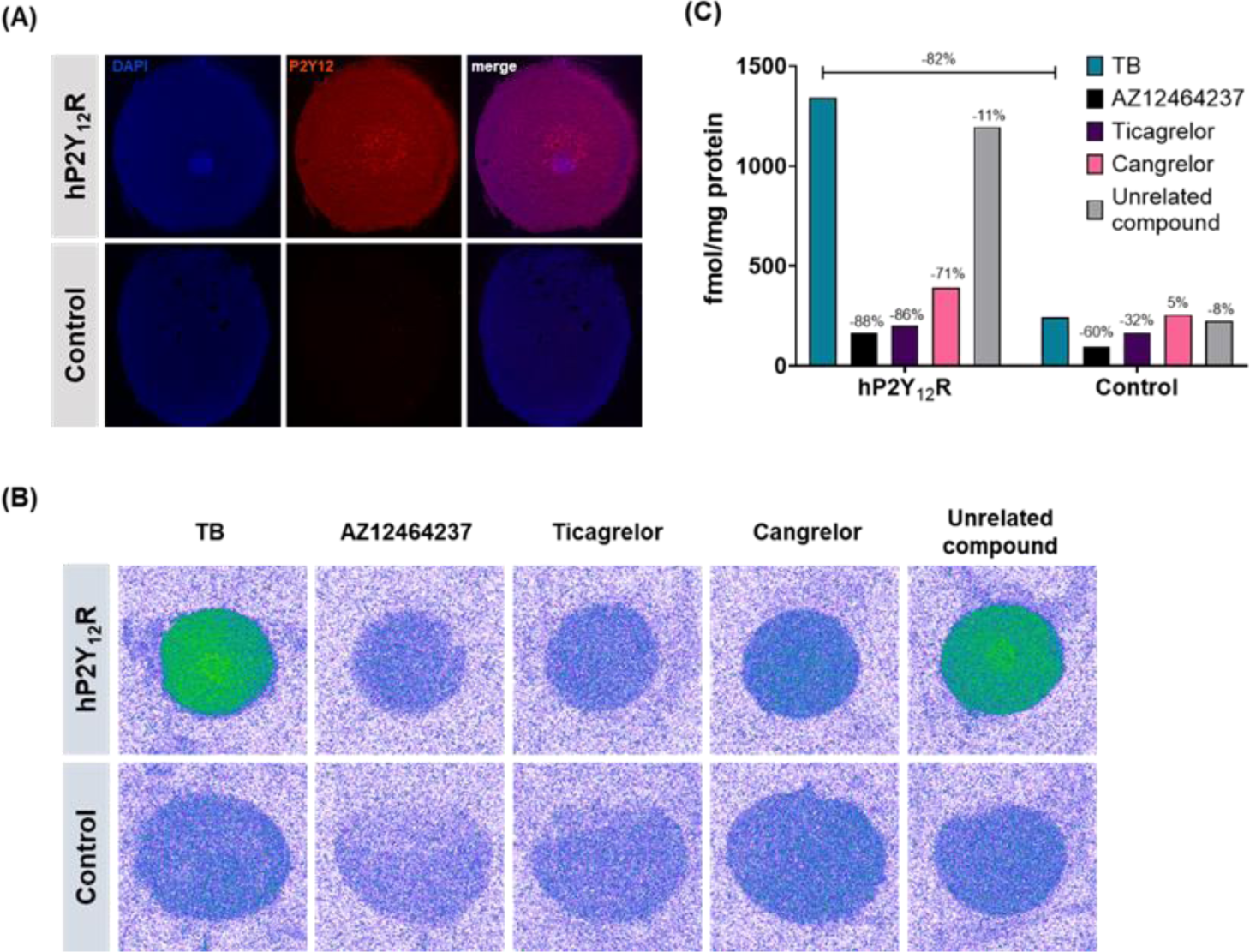
*In vitro* binding of [^3^H]AZ12464237 to frozen sections of cell pellets obtained from hP2Y_12_R-overexpressing and control CHO cells. **(A)** Expression of P2Y_12_R (red) on sections from P2Y_12_R-overexpressing and control CHO cells, scale bar: 50 μm. **(B)** TB of [^3^H]AZ12464237 (10 nM) and NSB in presence of AZ12464237, ticagrelor, cangrelor and a structurally unrelated compound (all at a 10 μM concentration) to P2Y_12_R overexpressing CHO cell pellets (top row) and control CHO cell pellets (bottom row). **(C)** Quantification of TB and NSB (expressed in fmol per mg of protein) to the P2Y_12_R overexpressing and control CHO sections. Representative graph of one experiment (experiment replicated three times).

With the promising results from the proof of concept autoradiography on sections from CHO cell pellets, the ability of [^3^H]AZ12464237 to detect the P2Y_12_R *in situ* was analyzed on brain tissue sections. Tracer binding was assessed on brain tissue derived from WT and P2Y_12_R knock-out (KO) mice. Immunolabeling against P2Y_12_R confirmed the expression of P2Y_12_R in the brain of WT mice by microglial cells (positive cells for Iba1), and verified that P2Y_12_R expression was completely absent in the brain sections from KO mice (Figure 8A). Binding of [^3^H]AZ12464237 to WT mouse brain sections could successfully be blocked by both unlabeled AZ12464237 (78% decrease), ticagrelor (65% decrease) and cangrelor (73% decrease) (Figure 8B and C). In addition, incubation of the radioligand in the presence of an unrelated compound did not result in any significant decrease in [^3^H]AZ12464237 binding, which supports the binding specificity of [^3^H]AZ12464237 to the P2Y_12_R. TB on brain sections derived from KO mice was lowered 55% compared to WT mouse brain sections. Interestingly, the binding levels in KO mouse brain sections was higher compared to the binding levels obtained by pharmacological blockade with ticagrelor or cangrelor. When co-incubating the radioligand with different blockers in the P2Y_12_R KO mouse brain sections, a further decrease of [^3^H]AZ12464237 was observed, in particular for cangrelor. These findings suggest the possibility of off-target binding by both AZ12464237 and cangrelor.

**Figure 8.**
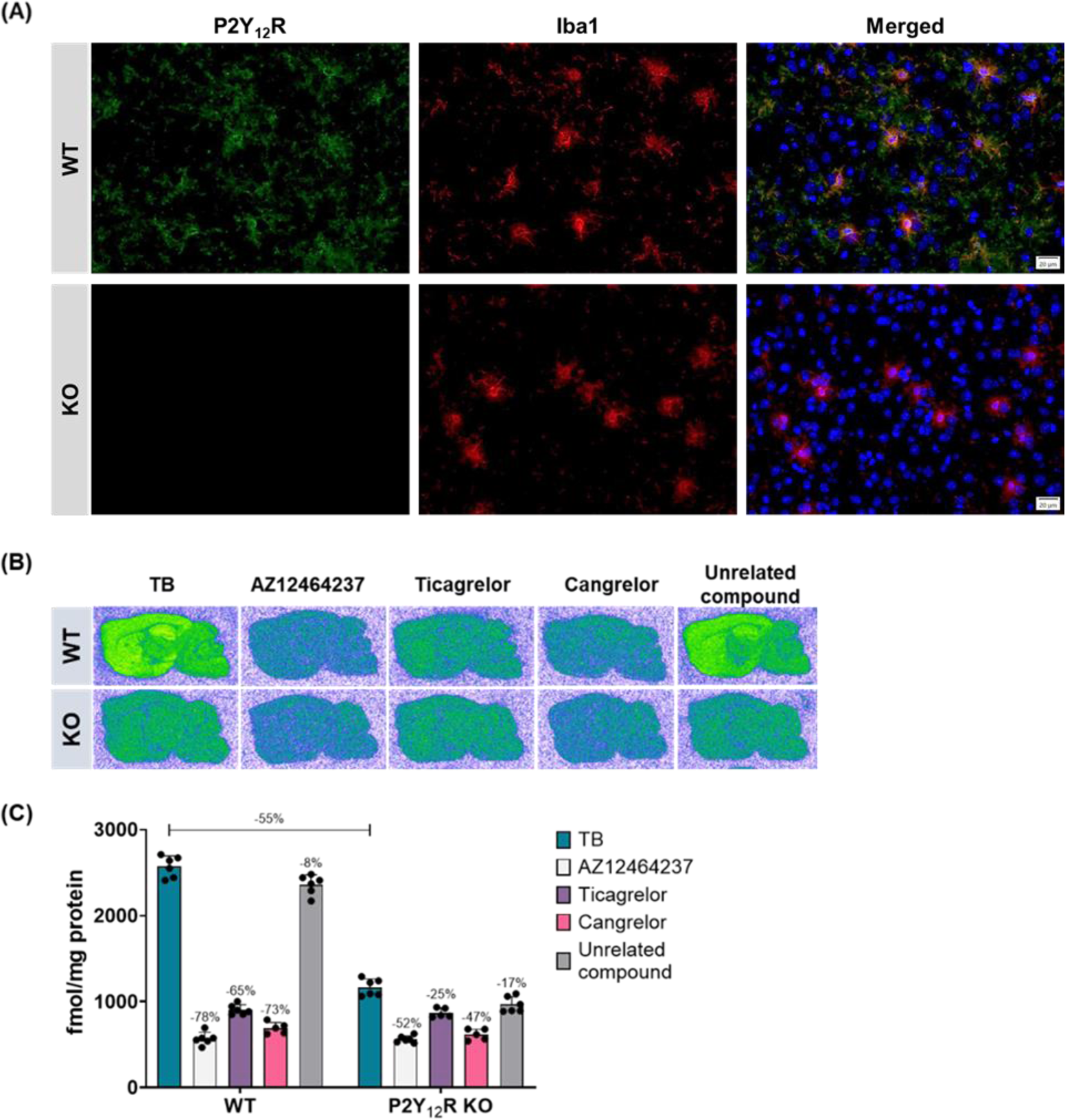
*In vitro* autoradiography with [^3^H]AZ12464237 on WT and P2Y_12_R KO mouse tissue. **(A)** Representative images of the expression of P2Y_12_R (green) and Iba1 (red) on the cortex of WT (top) and P2Y_12_R KO mouse tissue (bottom), scale bar: 20 μm. **(B)** Representative autoradiograms of [^3^H]AZ12464237 binding (10 nM) in the absence (TB) or in the presence of AZ12464237, ticagrelor, cangrelor or a structurally unrelated compound (all at a 10 μM concentration) on the sections. **(C)** Representative quantification of TB and NSB on WT and KO tissue (experiment replicated three times).

In order to further validate the selectivity of [^3^H]AZ12464237 and identify potential off-targets observed on the P2Y_12_R KO brain sections, a selectivity screening was performed. The non-radiolabeled AZ12464237 was screened at a concentration of 10 µM against a representative panel of 50 targets provided by Cerep.(34) In this screening two potential off-targets were identified: AZ12464237 displayed a high percentage of inhibition (95%, enzymatic activity assay) of glycogen synthase kinase 3 α (GSK3α), and a high displacement (84%, competition binding assay) at peroxisome proliferator-activated receptor γ (PPARγ). Additional experiments were carried out to investigate the dose response of AZ12464237 toward these two targets, which revealed an *IC_50_* value of 266 nM for GSK3α and a *K_i_* of 355 nM for PPARγ. With this information in hand, we repeated the autoradiography studies and included also reported ligands (Figure 9A) for the two identified off-targets namely, staurosporine (a broad-spectrum kinase inhibitor, including GSK3α) and rosiglitazone (PPARγ agonist). As shown in Figure 9B and C, co-incubation with rosiglitazone did not appear to have a significant effect on the binding of [^3^H]AZ12464237, neither on WT or P2Y_12_R KO tissues. Yet, co-incubation with staurosporine decreased the binding in both cases. Interestingly, the binding levels were similar when blocking with AZ12464237 (41% decrease) and staurosporine (48% decrease) in the P2Y_12_R KO mice, which further supports the hypothesis that GSK3α is the observed off-target on brain tissues.

**Figure 9.**
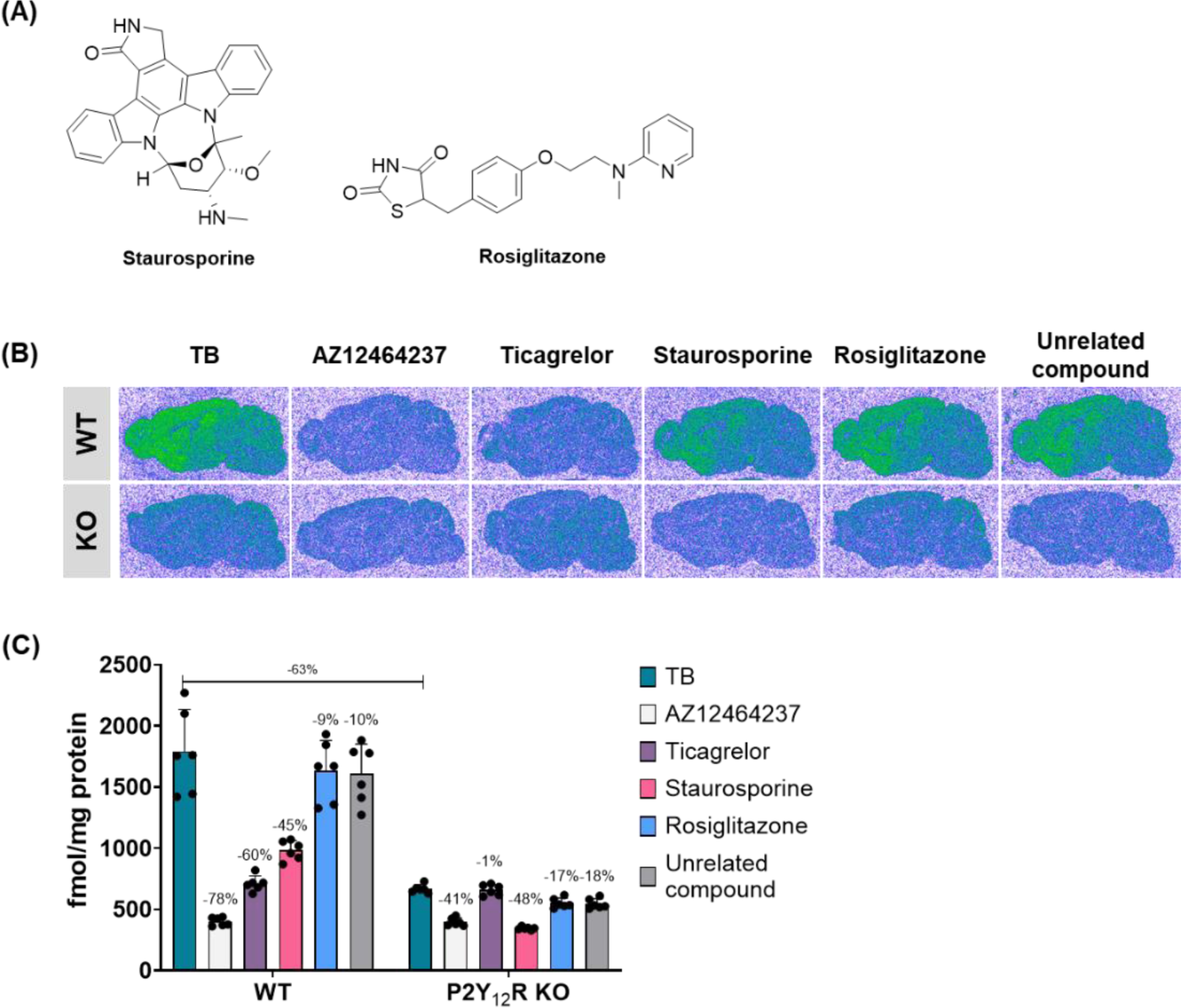
*In vitro* autoradiography with [^3^H]AZ12464237 on WT and P2Y_12_R KO mouse tissue. **(A)** Chemical structures of staurosporine and rosiglitazone. **(B)** Representative autoradiograms of [^3^H]AZ12464237 binding in the absence (TB) or in the presence of AZ12464237, ticagrelor, staurosporine, rosiglitazone or a structurally unrelated compound (all at a 10 μM concentration) on the sections. **(C)** Representative quantification of TB and NSB on WT and KO tissue (experiment replicated three times).

## 4. Discussion

The present study builds on the growing interest to study P2Y_12_R-mediated actions, as well as microglia dynamics in the CNS, and the merging need of an easily accessible pharmacological tool for this purpose.(17,35) Such a tool compound could be used in *in vitro* screening assays to identify novel brain-penetrant compounds that target the receptor in the CNS, as well as to further investigate P2Y_12_R in different cell types and brain tissues. Herein, we report the pharmacological characterization of [^3^H]AZ12464237, a novel radioligand based on a high-affinity P2Y_12_R antagonist.(23) In comparison to previously reported radioligands for the P2Y_12_R, [^3^H]AZ12464237 is based on a non-nucleoside scaffold and can be synthesized via a straightforward Pd-mediated tritiation in good yield and high RCP. Some of the earlier radioligands based on nucleosides have been challenging to synthesize due to their chemical complexity, making them either expensive or commercially unavailable.(17) *In vitro* the novel radioligand showed strong affinity (*K_d_* = 3.12 ± 0.70 nM) and saturable binding to the P2Y_12_R (Figure 4). The binding kinetics of [^3^H]AZ12464237 were rapid with a fast association and equilibrium reached within the first 10 min. The dissociation experiments revealed a very short target residence time (1.10 ± 0.22 min at room temperature). For comparison, ^3^H-labeled ticagrelor has been reported to have slower receptor kinetics with a residence time of approximately 19 min.(19) Moreover, our initial characterization studies confirmed that [^3^H]AZ12464237 specifically binds to hP2Y_12_R-HEK membranes, but not to membranes from HEK cells transfected with the P2X_7_R (Figure 3F). Since P2X_7_R is a recognized biomarker for the pro-inflammatory state of microglia (M1), which is the opposite state compared to the one linked to P2Y_12_R (M2), the lack of [^3^H]AZ12464237 binding to P2X_7_R is an encouraging result.(15,24) This suggests that the radioligand can be used alongside a selective P2X_7_R radioligand (*e.g.*, [^3^H]A-740003 and [^3^H]SMW139) to explore the relationship between microglial functional states at different stages of neuroinflammatory diseases *in vitro*.(15,36)

Among the evaluated P2Y_12_R antagonists, AZD1283 showed the greatest potency against [^3^H]AZ12464237 binding (*K_i_* = 2.66 ± 0.60 nM), followed by elinogrel (*K_i_* = 7.00 ± 0.54 nM). These ligands also have the highest structural similarity to the radioligand. The overall rank order of potency was as follows; AZD1283 > elinogrel > 2MeSADP ∼ ticagrelor > PSB0739. Our binding data is what one would expect from competitive studies against [^3^H]AZ12464237 for these compounds based on previous pharmacological data and structural information.(5–7,19,37) The rank order based on previously reported affinity data (Table 3) when competing with nucleoside-based radioligands can be summarized accordingly; 2MeSADP > ticagrelor > elinogrel ∼ AZD1283 ∼ PSB0739. As expected, when using a nucleoside-based radioligand the order will be reversed with competitors more similar to the used radioligand in the top of the ranking. Interestingly, the affinity of PSB0739 is ranked at the same place for both types of radioligands with similar *K_i_* values; 25.6 ± 3.05 nM against [^3^H]AZ12464237 and 25 ± 3.2 nM against the nucleotide derivative [^3^H]PSB0413. Solved crystal structures of the P2Y_12_R in complex with AZD1283, 2MeSADP or 2MeSATP, along with insights from molecular modeling and dynamics have revealed that non-nucleoside ligands bind to the receptor in a very different mode compared to the nucleotide derivatives. The binding site are only partially overlapping.(5–7) Additionally, there is a significant difference in receptor conformation when binding nucleotide derivatives versus uncharged, nucleoside-mimicking compounds, such as ticagrelor.

The selectivity of [^3^H]AZ12464237 binding was verified by performing competitive binding experiments with antagonists designed to target other receptors in the P2Y family (Table 3; Figure 5). In literature, the P2Y_13_R ligand MRS2211 is reported to have a > 20-fold selectivity toward P2Y_13_R over P2Y_1_R and P2Y_12_R, whereas, PPTN·HCl is described to be a highly selective P2Y_14_R antagonist (*K_b_* = 434 pM) with > 10 000-fold selectivity for P2Y_14_R over other P2Y receptors.(32,33) To our knowledge, no affinity data for MRS2578 toward the P2Y_12_R is reported in the literature. However, it is a highly selective P2Y_6_R antagonist (*IC_50_* = 37 ± 16 nM) over P2Y_1_R, P2Y_2_R, P2Y_4_R and P2Y_11_R (*IC_50_* >10 μM for all of them).(38) MRS2211 proved to be the most effective competitor against [^3^H]AZ12464237 binding, with a *K_i_* of 3.24 ± 1.85 µM, followed by PPTN·HCl and lastly MRS2578. This result could be explained by the fact that P2Y_13_R and P2Y_12_R share the same endogenous agonist, ADP, and the ligands target a somewhat similar binding site.

In order to prepare for future translational studies, we also investigated the use of [^3^H]AZ12464237 to label rP2Y_12_R, expressed in HEK cells, as well as the P2Y_12_R in brain tissue from mice in autoradiography studies. Even though the rodent rP2Y_12_R and mP2Y_12_R have a high amino acid homology with the human receptor (85% and 87% homology for rat and mouse protein respectively), species variation of GPCR targets are well-known to impact receptor-ligand interactions.(39,40) Consequently, the minor amino acid differences between human and rodent P2Y_12_Rs might still impact the binding of novel compounds, including [^3^H]AZ12464237. Yet, [^3^H]AZ12464237 binding on membrane preparations with the rP2Y_12_R also resulted in high affinity and saturable labeling of the receptor. The *K_d_* and *B_max_* values for [^3^H]AZ12464237 for rP2Y_12_R were higher compared to the values obtained for the hP2Y_12_R. However, this trend was also observed in the competitive [^3^H]AZ12464237 displacement studies with ticagrelor and elinogrel. Subsequent *in vitro* autoradiography experiments on mouse brain tissue further confirmed the species translatability of the radioligand (Figure 8). [^3^H]AZ12464237 could detect the P2Y_12_R on brain tissues from wild-type mice and the binding could successfully be blocked with both ticagrelor, cangrelor and the non-radiolabeled analog of AZ12464237. The SB (based on ticagrelor blocking) was lower on brain tissues, compared to the transfected cell lines, 65% versus ∼90% at concentrations around the *K_d_* values. However, in P2Y_12_R KO mice we observed a rather high level of [^3^H]AZ12464237 binding (decreased 55% compared to the WT mice tissue), which could also be blocked with AZ12464237 and cangrelor, but not significantly with ticagrelor. Suspecting off-target binding as the cause of the observed results, a comprehensive selectivity screening was performed at Cerep. This screening identified two potential off-targets for AZ12464237: GSK3α and PPARγ. To further investigate these, we repeated the autoradiography experiments, this time including GSK3α and PPARγ ligands as blockers (Figure 9). We were able to attribute the off-target binding to GSK3α, as co-incubation with AZ12464237 and the GSK3α inhibitor staurosporine reduced radioligand binding to similar levels in P2Y_12_R KO mice. However, we do not believe this minor off-target binding will interfere with the use of the radioligand in *in vitro* studies, as the affinity of AZ12464237 for P2Y_12_R is nearly 90 times higher than its affinity for GSK3α.

In conclusion, we report the radiosynthesis and *in vitro* binding characterization of the non-nucleoside P2Y_12_R antagonist [^3^H]AZ12464237. The ^3^H-labeled ligand exhibits high affinity binding to P2Y_12_R with rapid kinetics, and demonstrated binding specificity. Moreover, [^3^H]AZ12464237 is the first non-nucleoside radioligand for the P2Y_12_R. Its straightforward synthesis makes it an easily accessible tool compound for *in vitro* studies including screening assays and autoradiography experiments to uncover the exact role of the P2Y_12_R in the brain, particularly in the context of microglia activation in neuroinflammatory diseases.

## CRediT authorship contribution statement

**E. Johanna L. Stéen:** Conceptualization, Investigation, Methodology, Formal analysis, Writing – original draft, Writing – review & editing, Funding acquisition. **Efthalia-Natalia Tousiaki:** Investigation, Methodology, Formal analysis, Writing – review & editing. **Lee Kingston:** Investigation, Formal analysis, Writing – review & editing. **Berend van der Wildt:** Investigation, Methodology, Writing – review & editing, Funding acquisition. **Nikolett Lénárt:** Investigation, Methodology, Writing – review & editing. **Wissam Beaino:** Investigation, Methodology, Formal analysis, Writing – review & editing, Supervision. **Mariska Verlaan:** Investigation, Methodology, Writing – review & editing. **Barbara Zarzycka:** Resources, Formal analysis, Writing – review & editing. **Bastian Zinnhardt:** Formal analysis, Writing – review & editing. Supervision, Project administration. **Ádám Dénes:** Resources, Formal analysis, Writing – review & editing**. Luca Gobbi:** Resources, Formal analysis, Writing – review & editing, Funding acquisition, Supervision. **Iwan J.P. de Esch:** Resources, Writing – review & editing, Funding acquisition. **Charles S. Elmore:** Formal analysis, Writing – review & editing, Project administration. **Albert D. Windhorst:** Conceptualization, Formal analysis, Writing – review & editing, Project administration, Funding acquisition. **Michael Honer:** Conceptualization, Resources, Formal analysis, Writing – review & editing, Project administration. **Rob Leurs:** Conceptualization, Resources, Formal analysis, Writing – original draft, Writing – review & editing.

## Abbreviations

ADP: adenosine-5’-diphosphate
ATP: adenosine-5’-triphosphate
AZD1283: Ethyl 5-cyano-2- methyl-6-[4-[[[(phenylmethyl)sulfonyl]amino]carbonyl]-1-piperidinyl]-3-pyridinecarboxylate
*B_max_*: maximal binding at equilibrium
Cangrelor: *N*-[2-(methylthio)ethyl]-2-[(3,3,3- trifluoropropyl)thio]adenosine-5’-*O*-(β,γ-dichloromethylene)triphosphate
CHO: Chinese hamster ovary
CNS: central nervous system
Cys: cysteine
DMSO: dimethylsulfoxide
Elinogrel: 5-chloro- *N*-[[[4-[6-Fluoro-1,4-dihydro-7-(methylamino)-2,4-dioxo-3(2*H*)- quinazolinyl]phenyl]amino]carbonyl]-2-thiophenesulfonamide
GPCR: G protein-coupled receptor
HEK 293: human embryonic kidney 293
HEPES: *N*-(2-hydroxyethyl)piperazine-*N*’-(2- ethanesulfonic acid)
*IC_50_*: half maximal inhibitory concentration
*K_d_*: equilibrium dissociation constant
*K_i_*: equilibrium inhibitory constant
KO: knock-out
*K_off_*: dissociation rate constant
*K_on_*: association rate constant
MRS221: 2-[(2-chloro-5-nitrophenyl)azo]-5-hydroxy-6-methyl-3- [(phosphonooxy)methyl]-4-pyridinecarboxaldehyde disodium salt
MRS2578: *N*,*N’’*-1,4- butanediylbis[*N’*-(3-isothiocyanatophenyl)thiourea
nM: nanomolar
NSB: non-specific binding
P2Y_12_R: purinergic 2Y type 12 receptor
P2X_7_R: purinergic 2X type 7 receptor
PET: positron emission tomography
PPTN·HCl: 4-[4-(4-piperidinyl)phenyl]-7-[4-(trifluoromethyl)phenyl]-2- naphthalenecarboxylic acid hydrochloride
Rosiglitazone: 5-[[4-[2-(methyl-2- pyridinylamino)ethoxy]phenyl]methyl]-2,4-thiazolidinedione
SB: specific binding
SEM: standard error of the mean
Staurosporine: [9S-(9α,10β,11β,13α)]-2,3,10,11,12,13-Hexahydro-10-methoxy- 9-methyl-11-(methylamino)-9,13-epoxy-1H,9H-diindolo[1,2,3-gh:3’,2’,1’-lm]pyrrolo[3,4- j][1,7]benzodiazonin-1-one
TB: total binding
TEA: triethylamine
Tris: tris(hydroxymethyl)aminomethane
UDP: uridine diphosphate
UDPG: uridine diphosphate-sugar derivatives (galactose/glucose)
UTP: uridine triphosphate
WT: wild-type

## Acknowledgements

The authors would like to thank Ileana Guzzetti at AstraZeneca for technical assistance with the ^3^H NMR studies. This project received funding from the Michael J. Fox foundation under grant agreement No. MJFF-022453 (E.J.L.S, L.G., I.J.P.E, A.D.W., M.H., and B.Z.). E.J.L.S received funding from the European Union’s Horizon 2020 research and innovation programme as a Marie Sklodowska-Curie Individual Fellowship under grant agreement No. 892572: Molecular Imaging of Microglia (MIM). B.v.d.W was funded by the Dutch Research Council (NWO) with a NWO-VENI grant under grant agreement No. VI.Veni.192.229.

## Data availability

The authors declare that all data supporting the findings described are available within the paper. Raw data are available from the corresponding author upon reasonable request.

